# Disruption of sRNA Function Using Synthetic Arginine Rich Motif Peptides

**DOI:** 10.64898/2026.08.19.745773

**Authors:** Edwin E. Ortiz, Arada J. Batresian, Jezriel D. Punzalan, Andrea Gutierrez Garcia, Briet Bjornsson, Ivan Khoroz, Ravinder Abrol, Melissa K. Takahashi

## Abstract

Small RNAs (sRNAs) regulate the expression of many genes including those involved in antibiotic resistance and bacterial virulence, making them potential therapeutic targets. A molecule that binds an sRNA could interfere with its ability to bind its target mRNA and disrupt the regulation mechanism. Randomization and screening of natural arginine rich motif (ARM) peptides led to peptides capable of interfering with the sRNA MicF’s ability to regulate *ompF* in *Escherichia coli*. Molecular dynamics simulations suggested that this effect was not a result of a direct disruption of the MicF-*ompF* interaction. Instead, the peptides interfere with binding of the chaperone Hfq, which is required for MicF-mediated regulation. Subsequent testing demonstrated peptide specificity for MicF over two other Hfq scaffolds and the ability to disrupt regulation of two additional MicF targets. Together, these findings support the use of synthetic ARMs as a potential tool for modulating sRNA function in bacteria.

## Introduction

Small RNAs (sRNAs) are regulatory molecules that allow bacteria to rapidly respond to changes in their environment^1,2^. sRNAs are known to regulate the expression of many genes including those involved in antibiotic resistance and virulence mechanisms such as outer membrane proteins, motility, biofilm formation, and quorum sensing^2–5^. Given the continued concern surrounding antibiotic-resistant pathogens, sRNAs could serve as new drug targets^6^. sRNAs regulate gene expression post transcriptionally by directly binding to mRNAs, often with the help of a protein chaperone^2,3^. Binding between sRNA and mRNA changes the accessibility of the ribosome binding site or RNase-based degradation of the mRNA. Therefore, a molecule that binds or sequesters the sRNA would prevent regulation of the mRNA and disrupt the antibiotic resistance or virulence mechanism of the bacteria.

Prior work by us and others have demonstrated the feasibility of targeting sRNAs with the use of antisense oligonucleotides (ASOs) that directly base pair to the sRNA^7,8^. However, because of the length requirements associated with the delivery of some ASOs into bacteria^9,10^, multiple molecules may be required to sequester a given sRNA^8^. As an alternative approach for targeting sRNAs, we explored the use of peptides, specifically arginine rich motif (ARM) peptides. ARMs are characterized by an abundance of arginine residues that facilitate binding to a specific RNA. ARM-RNA binding results in various functions in nature including regulation of antitermination, mRNA transport, and splicing^11,12^. Studies have shown that ARMs bind to their cognate RNAs with high specificity^13,14^ and their binding affinity can be improved by mutating a few amino acids within the ARM^15,16^. In other work, a synthetic ARM that binds to the human immunodeficiency virus (HIV) Rev response element RNA (RRE) was identified from a combinatorial library of peptides^17^. These results led us to investigate if an ARM could be designed to bind an sRNA.

In this work, we present a strategy to randomize the sequence of ARMs found in nature and screen for the ability to interfere with sRNA-mRNA binding. To demonstrate the strategy, we target the sRNA MicF, which we have previously designed ASOs to inhibit^8^. MicF is a 93-nucleotide sRNA, that is induced under several conditions including high osmolarity, oxidative stress, and the presence of antibiotics^18–24^. One of the mRNAs that MicF regulates is that of the outer membrane protein OmpF, which in addition to nutrients, is an entry way for some antibiotics^25,26^. Deletion or inhibition of MicF leads to increased susceptibility to those antibiotics^8,23^. We generated libraries of ARMs, screened for peptides that interfered with MicF-*ompF* binding, and identified two peptides that increased susceptibility of *Escherichia coli* to cephalothin. Molecular dynamics (MD) simulations and supporting experiments suggest that the identified peptides bind to MicF in the same region that the sRNA chaperone Hfq binds. These results indicate the potential for using peptides to target sRNAs.

## Results and Discussion

### Workflow to Generate and Screen Potential MicF-Binding Peptides

To identify peptides that interfere with MicF-*ompF* binding, we utilized an experimental setup analogous to the one used to screen anti-MicF antisense RNAs^8^ (Figure 1A). The reporter was a construct that fused the 5’ untranslated region (UTR) and first 13 codons of *ompF* to the coding sequence of superfolder green fluorescent protein (sfgfp)^27^. The reporter plasmid was transformed into *E. coli* along with two additional plasmids containing MicF and a peptide or control. Each component was constitutively expressed using a medium strength J23118 promoter^28^ to minimize burden on cell growth. The anti-MicF 33-1 antisense RNA (asRNA) from Tsai et al.^8^ was used as a control to represent the maximum expected interference of MicF-*ompF* binding. Four ARMs were chosen as starting peptides: the N proteins from bacteriophages λ and P22, the Tat protein from bovine immunodeficiency virus (BIV), and the Rev protein from HIV. A preliminary test was performed to determine if any of the ARMs naturally interfered with MicF-*ompF* binding or the *ompF::sfgfp* reporter, which was not the case (Figure 1B).

**Figure 1.**
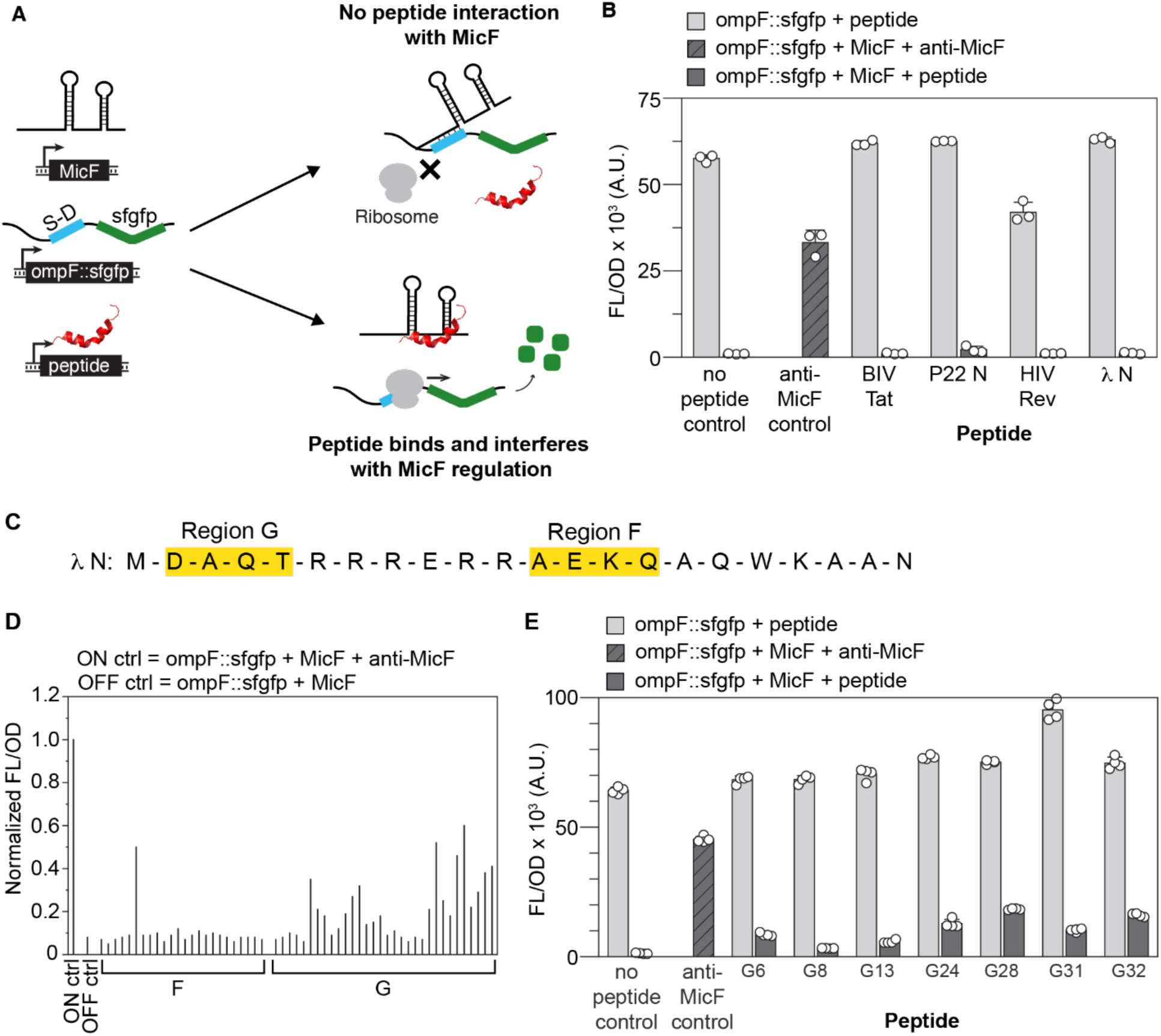
Peptide screening workflow. (A) Schematic of the peptide screening assay. Plasmids encoding constitutively expressed MicF, *ompF::sfgfp*, and peptide candidates are transformed into *E. coli*. Cells containing a peptide that interferes with MicF-*ompF* binding express SFGFP while cells containing peptides that do not interact with MicF do not express SFGFP. (B) Initial test to determine if any of the natural ARMs interfere with MicF-*ompF* binding. Bars show mean fluorescence/optical density (FL/OD). Error bars represent the standard deviations of three biological replicates, shown as open circles. (C) Amino acid (AA) sequence of the λ N peptide. Two regions of AA, F and G highlighted in yellow, were randomized independently via PCR (see Methods). (D) Individual colonies from the transformations depicted in (A) were grown in liquid culture. FL/OD for each individual culture was normalized to the ON ctrl. (E) Peptides from (D) that resulted in normalized FL/OD of at least 0.2 were sequenced. Those with clean, full-length peptide sequences were cloned into plasmids and tested in *E. coli* along with the *ompF::sfgfp* reporter and MicF. Bars show mean FL/OD. Error bars represent the standard deviations of four biological replicates, shown as open circles.

For each peptide, the amino acids chosen for randomization were those that were shown to be involved in the binding of the peptide to its native RNA target^29–31^ while leaving any arginines in place. The amino acids were grouped together in regions to be randomized (Figure 1C, S1A). Randomization was achieved through PCR by ordering primers with N’s in the positions of the selected amino acid codons. After PCR amplification and blunt-end ligation, cloned plasmids were transformed into competent cells containing the reporter and MicF plasmids. Colonies were screened visually for sfgfp fluorescence using a transilluminator. Visually green colonies were used to inoculate liquid cultures and measured for bulk fluorescence (FL) and optical density (OD). Although no green colonies were observed for the randomized P22-N, BIV-Tat, and HIV-Rev peptides, several were selected for measurement in liquid culture to determine if any increase in fluorescence could be measured (Figure S1B). The only potential hits from the initial screen came from the randomization of the λ N peptide (Figure 1D). Upon sequencing, it was noted that several of the screened colonies contained multiple peptide sequences and others had truncated peptides (e.g. the hit from the F region randomization). Only the full-length peptide sequences obtained from the screen were cloned into plasmids and tested again with the *ompF::sfgfp* reporter and MicF (Figure 1E). Several of the peptides demonstrated an ability to interfere with MicF-*ompF* binding although none as well as the anti-MicF asRNA control.

To improve the interaction between the peptides and MicF, the hits from the initial screen were used as templates for a second round of randomization. The G regions for each peptide were held constant while the F and an additional H region were randomized (Figure 2, S2-S4). For this round of randomization, the visual screen could not be used since all colonies were green. Instead, colonies were used to inoculate liquid cultures, and the FL/OD values were compared to the original peptide used for randomization. Between 180-356 colonies (peptides) were screened from each of the λ N G24, G28, G31, and G32 F and H randomizations. Colonies with normalized FL/OD values above a set threshold were sequenced and those with clean, full-length peptide sequences were cloned and tested again as above. Except for the λ N G24 peptide, the F and H randomizations yielded peptides that demonstrated greater interference with MicF-*ompF* binding than the original peptide (Figures 2B, S2).

**Figure 2.**
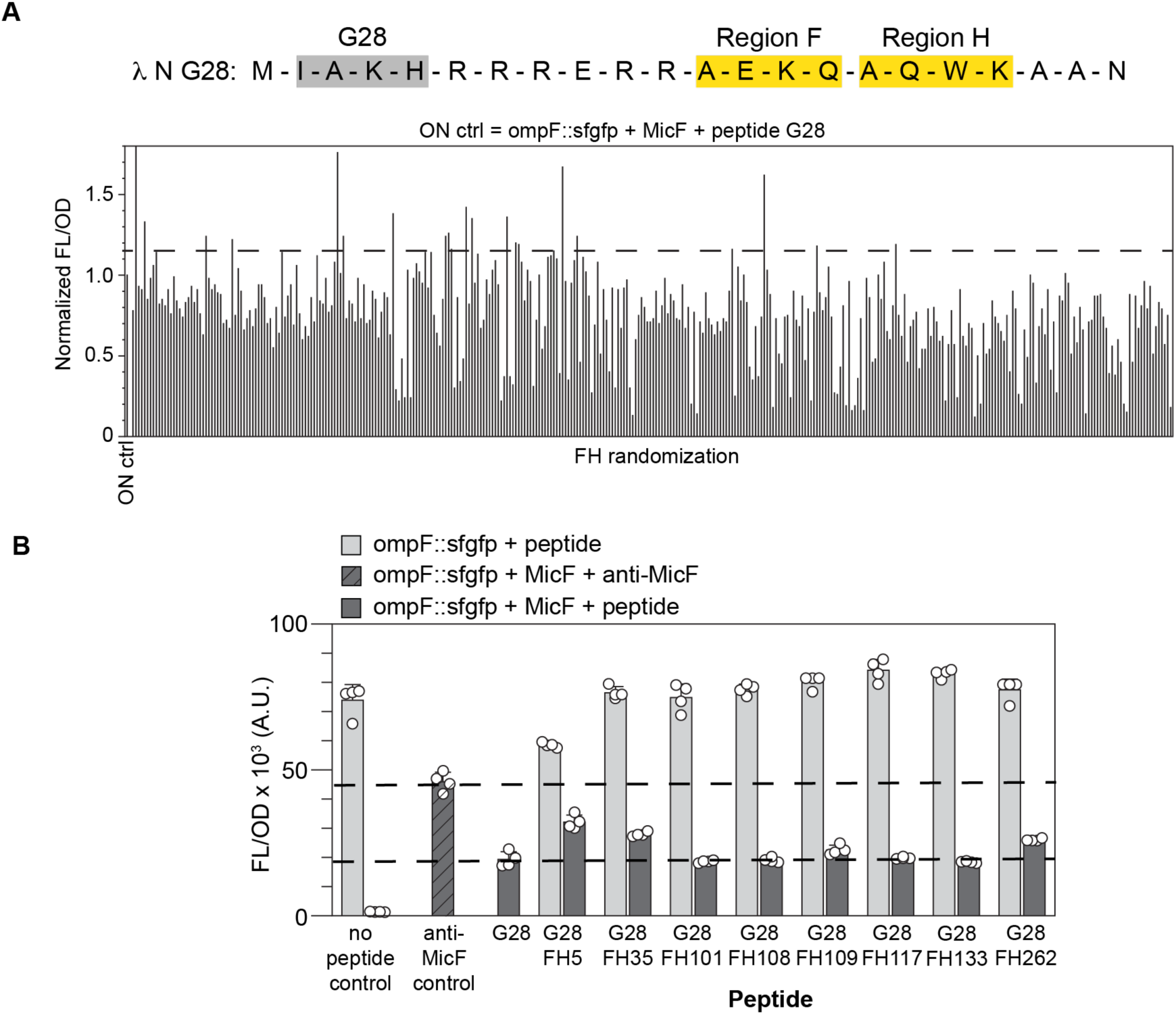
Randomization of peptide λ N G28. (A) Randomization screen. The AA sequence of the λ N G28 peptide is shown with the randomized regions F and H highlighted in yellow. Individual colonies from the randomization cloning were grown in liquid culture. FL/OD for each individual culture was normalized to the ON ctrl which was the *ompF::sfgfp* reporter with MicF and the λ N G28 peptide. Peptides above the threshold depicted by the dashed line were sequenced. (B) Peptides above the threshold in (A) with clean, full-length peptide sequences were cloned and tested in *E. coli* along with the *ompF::sfgfp* reporter and MicF. Bars show mean FL/OD. Error bars represent the standard deviations of four biological replicates, shown as open circles. Dashed lines are drawn for comparison at the FL/OD of the anti-MicF control (upper line) and λ N G28 peptide (lower line).

### Minimum Inhibitory Concentration (MIC) Testing of Peptides

Given that the best peptides from the randomization screens were only 57-70% as effective as the anti-MicF asRNA control (Figure 2B), we sought to determine if the peptides could still interfere with *ompF* production in *E. coli*. To do this, we cloned the λ N G28_FH35 and G28_FH262 peptides on a plasmid downstream of the inducible promoter pLux^32^. Although the λ N G28_FH5 peptide demonstrated greater interference with MicF-*ompF* binding, it also interfered with expression of the *ompF::sfgfp* reporter and therefore was not selected for further testing. *E. coli* MG1655 cells were transformed with the plasmid containing the G28_FH35, G28_FH262, or original λ N peptide as a control. Cephalothin broth microdilution MIC assays were performed on each strain with *N*-acyl homoserine lactone (AHL) added at the time of antibiotic addition to induce peptide production. We observed a decrease in the MIC in the presence of both peptides (Figure 3A). The reduction in MIC was comparable to that seen in an *E. coli* strain where MicF was deleted from the genome^23^ and *E. coli* in the presence of anti-MicF ASOs^8^.

**Figure 3.**
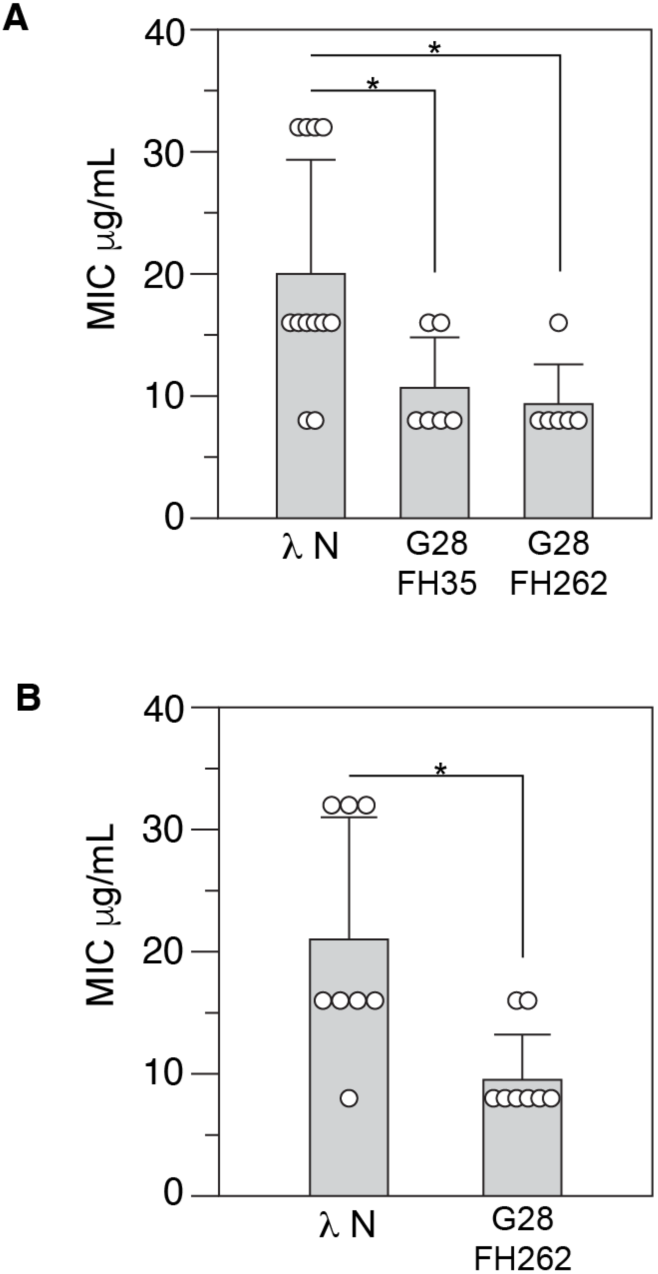
Evaluation of peptide effectiveness using MIC assay. (A) MIC for cephalothin for *E. coli* MG1655 expressing the λ N, G28_FH35, and G28_FH262 peptides. Statistical significance was assessed using an ANOVA followed by paired Welch’s t-test with Holm correction for multiple comparisons; *p* < 0.05 (*). (B) MIC for cephalothin for *E. coli* MG1655 conjugated with a plasmid carrying G28_FH262. Statistical significance was assessed using a Welch’s t-test; *p* < 0.05 (*).

The MIC result suggests that peptides could be an alternative to ASOs for interfering with sRNA-mRNA binding; however, there is a greater concern with delivering peptides into bacterial cells because of their potential cytotoxicity^33^. Two potential methods for peptide delivery that would alleviate this concern is the use of bacterial conjugation or bacteriophage to deliver genes to express the peptide specifically to bacterial cells^34–38^. Each method would require development beyond the scope of this work; however, as a proof-of-concept, we utilized a conjugation assay to deliver the λ N G28_FH262 peptide to *E. coli* MG1655 cells and performed MIC assays on the recipient cells for cephalothin. For conjugation, the p_Lux_-peptide sequence was cloned into a plasmid with the RSF1010 oriT. *E. coli* WM3064 was used as a donor strain and presence of the plasmid containing the peptide was confirmed via colony PCR performed on recipient MG1655 cells (Figure S5). MIC assays were performed as before comparing a strain with the λ N G28_FH262 peptide to one with the original λ N peptide. Again, we observed a decrease in the MIC for cephalothin (Figure 3B).

### Molecular Dynamics Simulations Suggest Peptides Interfere with Hfq binding

To investigate the interactions between the peptides and MicF, MD simulations were run on peptide-MicF complexes for the original λ N, G8, G28_FH35, and G28_FH262 peptides (Figure 4). The λ N and G8 peptides were selected as negative controls since they did not interfere with MicF’s regulation of OmpF in our reporter assay (Figure 1B, 1E). Our initial hypothesis was that peptides capable of interfering with MicF-*ompF* binding would create stable complexes with MicF; while peptides incapable of interfering with MicF would not. However, a free energy (ΔG) analysis indicated that all complexes were stable (Figure S6). As expected, the arginines within the peptides contribute significantly to the stable complexes formed with MicF indicated by an analysis of the electrostatic energy contributions from the individual amino acid residues (Tables S5-S8). Analysis of the interactions between individual amino acids within the peptides and the nucleotides of MicF revealed a lack of interaction with the first 33 nucleotides of MicF, which are responsible for the direct binding to *ompF*^24^. Instead, the arginines in the G28_FH262 peptide interact with MicF further downstream along the AU motif and poly U tail that Hfq is predicted to bind to^39^ (MicF residues 60-63 and 86-93) while the G28_FH35 peptide mostly interacts with the AU motif (Figure 4, Tables S11-S12). For the λ N and G8 peptides, the arginines and two additional lysines that contribute to the interaction with MicF (residues 14 and 19) interact more widely across MicF rather than concentrated in the predicted Hfq binding regions (Tables S9-S10). This suggested that rather than direct interference of MicF-*ompF* binding, the peptides interfere with MicF’s binding to Hfq, which is known to be required for regulation by MicF^27,40,41^.

**Figure 4.**
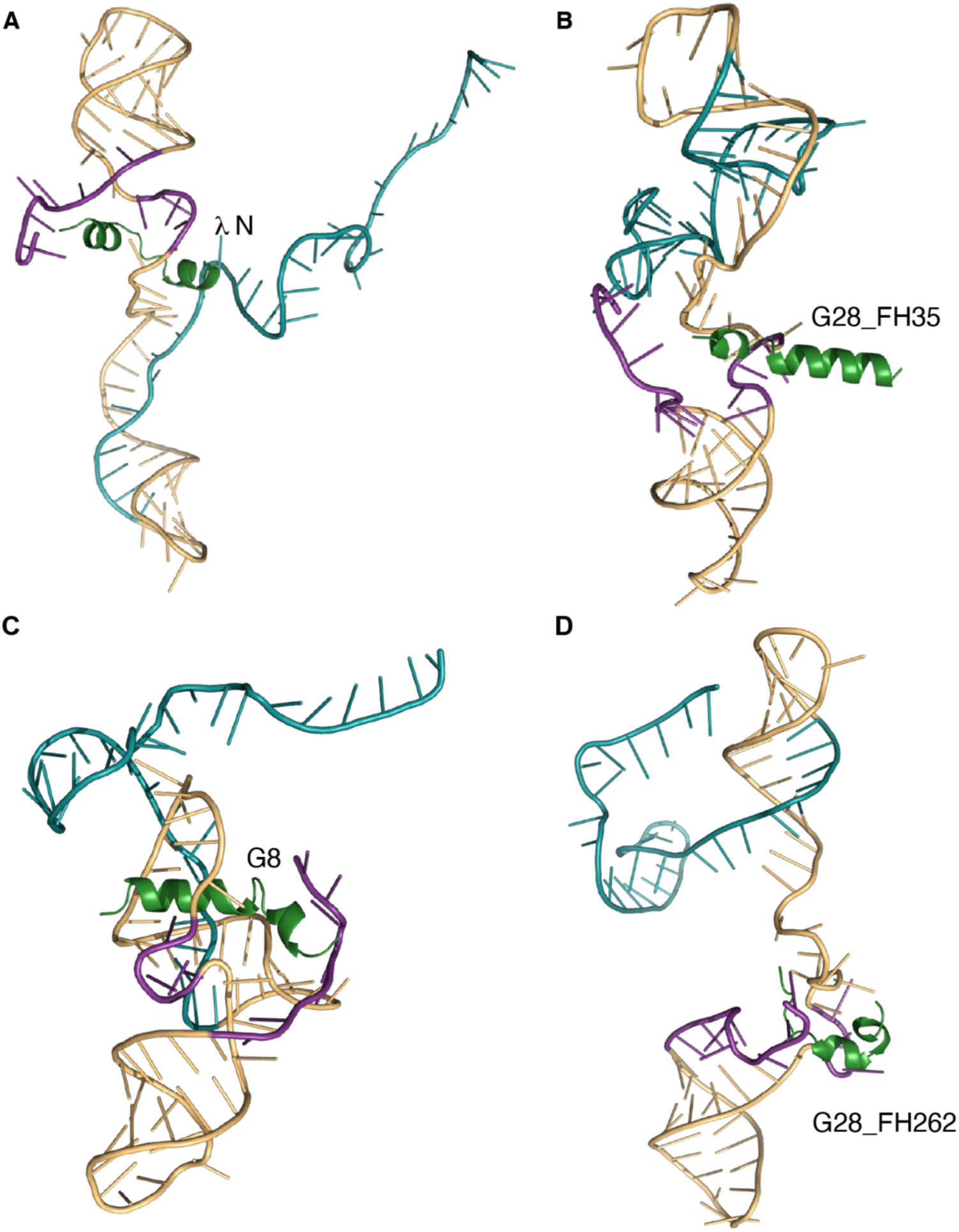
Last snapshots of MicF-peptide complexes from molecular dynamics (MD) simulations. (A) λ N-MicF. (B) G28_FH35-MicF. (C) G8-MicF. (D) G28_FH262-MicF. Images rendered with PyMOL. Peptides are colored green. Predicted Hfq binding regions of MicF are colored purple. The first 33 nucleotides of MicF, which are required for binding to *ompF* are colored blue.

To support the simulation findings, several additional tests were performed. If the peptides prevent Hfq from binding to MicF, they should also interfere with MicF’s ability to regulate its other targets. To test this, sfgfp fusions with two other MicF targets (Lrp and ObgE) were utilized^41^. Both λ N G28_FH35 and G28_FH262 demonstrated the ability to interfere with MicF regulation of *lrp::sfgfp* and *obgE::sfgfp* (Figure 5 A-B). Next, the peptides were tested with a MicF construct that took the first 13 nucleotides of MicF and fused it to the Hfq scaffold of the SgrS sRNA. The MicF(1-13) SS construct is known to repress *ompF::sfgfp* ^41^ and the SgrS Hfq scaffold has a distinct sequence from the MicF Hfq scaffold. Neither λ N G28_FH35 nor G28_FH262 interfered with the ability of MicF(1-13) SS to repress *ompF::sfgfp* (Figure 5C). Finally, OmpF expression is also repressed by the sRNA RybB^42^. RybB also has a Hfq-binding scaffold distinct from MicF and again, neither peptide interfered with the ability of RybB to repress *ompF::sfgfp* (Figure 5D). Together, these results support the notion that the peptides bind to MicF in the region that Hfq does, thus preventing Hfq binding and interfering with MicF’s ability to regulate its targets.

**Figure 5.**
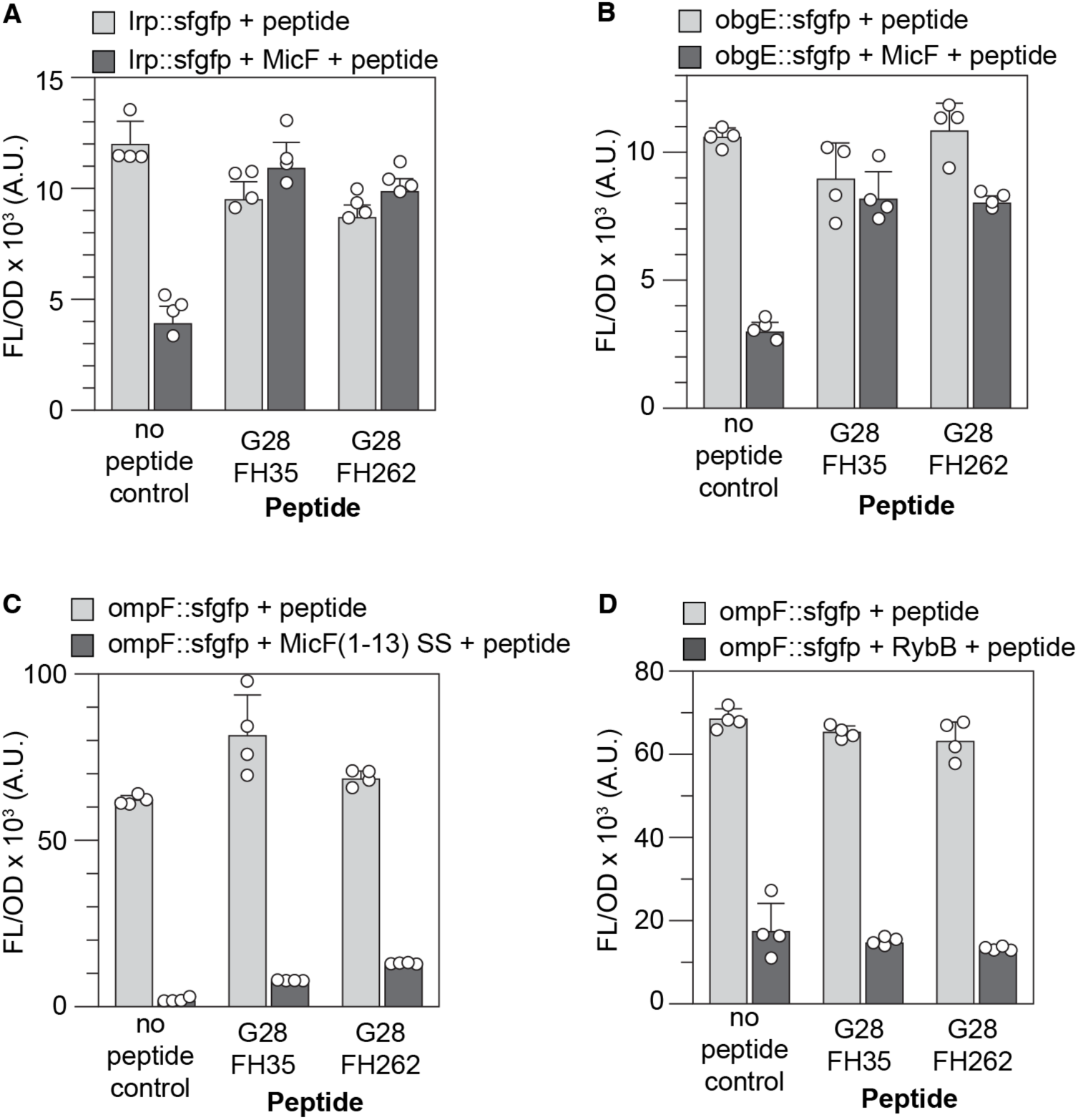
Tests to support peptides G28_FH35 and G28_FH262 binding to MicF in the Hfq binding scaffold. (A) *E. coli* transformed with plasmids expressing *lrp::sfgfp*, MicF, and peptides. (B) *E. coli* transformed with plasmids expressing *obgE::sfgfp*, MicF, and peptides. (C) *E. coli* transformed with plasmids expressing *ompF::sfgfp*, MicF(1-13) SS, and peptides. (D) *E. coli* transformed with plasmids expressing *ompF::sfgfp*, RybB, and peptides. Bars show mean FL/OD. Error bars represent the standard deviations of four biological replicates, shown as open circles.

## Conclusions

In this work, we demonstrated that ARMs can be designed to bind to the sRNA MicF and inhibit MicF’s ability to regulate its mRNA targets. The peptides were designed by randomizing the amino acid sequences of ARMs found in nature and then screening the peptides for the ability to interfere with MicF’s regulation of an *ompF::sfgfp* reporter. Our MD simulations and experiments suggest that the peptides are preventing Hfq from binding to MicF and thus disrupting MicF’s ability to bind to its mRNA targets. Although limited testing was performed, the peptides do show specificity for MicF over RybB and potentially SgrS.

The workflow we presented could easily be adapted to other sRNA-mRNA pairs. Several other methods have been developed to screen for RNA-protein interactions^17,43–46^. However, since the MD simulations suggest that other peptides make stable complexes with MicF, the use of the mRNA-sfgfp fusion reporter only identifies the peptides that interfere with regulation. Furthermore, the use of mRNA-protein reporter fusions is common practice in studying sRNA biology^40,47^. Regarding peptide design, there are two modifications that could increase the diversity of peptide sequences. Instead of starting with natural ARMs, Harada et al.^17^ constructed a library of 14-residue peptides consisting of arginine, serine, asparagine, or histidine at each position with a bias towards arginine; thus, creating a new ARM scaffold. Additionally, instead of using a nucleotide-based mutagenesis for the randomization, a codon-based mutagenesis approach would reduce any amino acid bias^16,48^.

sRNA-binding peptides add to the toolbox of molecules that can be designed to inhibit sRNA regulation that also include ASOs^7,8^ and sponge RNAs^49^. Given the prevalence of sRNA regulation in antibiotic resistance and virulence mechanisms, these molecules could improve the effectiveness of existing antibiotics or be used in an anti-virulence strategy that would enhance the immune system’s ability to clear bacterial pathogens^6,50–52^.

## Methods

### Strains and growth media

Three different *E. coli* strains were used throughout the study. NEB Turbo (New England Biolabs) was used for cloning and randomization protocols. MG1655 was used for MIC and bulk fluorescence testing. WM3064 was used as the donor strain for conjugation and selected for using 0.3 mM diaminopimelic acid (DAP). Strains were grown in LB broth, LB agar (1.5%), or MOPS EZ Rich Defined Media (Teknova M2105). The following antibiotics were used for plasmid selection: carbenicillin (carb) 100 μg/ml, chloramphenicol (cm) 34 μg/ml, and kanamycin (kan) 50 μg/ml.

### Plasmids

All plasmids used in this study are listed in Table S4 with key DNA sequences listed in Table S1.

### Construction and screening of plasmids with randomized ARM sequences

Randomized ARMs were generated by inverse PCR cloning. Plasmids containing the base peptide (e.g. λ N) were used as the template DNA for the PCR with primers used listed in Table S2. Blunt-end ligated DNA was transformed into chemically competent *E. coli* Turbo cells that contained the *ompF::sfgfp* reporter (MKT172) and MicF (MKT173). Transformed cells were plated onto LB-Agar with carb, cm, and kan and incubated overnight at 37°C. Individual colonies were used to inoculate 300 μl of LB with the corresponding antibiotics in a 2 ml 96-well block (Costar 3961). Three of the 96-wells were used as media blanks and an additional three were inoculated with colonies containing MKT172, MKT173, and the antiMicF(33-1) (MKT174) or template peptide plasmid. The block was sealed with a breathe-easier membrane (USA Scientific 9126-2100) and grown for 17 hours overnight at 37°C while shaking at 100 rpm (Labnet Vortemp 56). Four microliters of the overnight culture were added to 296 μl of LB in a new 96-well block and incubated for 3 hours under the same conditions as above. After incubation, 50 μl of sterile water and 50 μl of each culture was transferred to a 96-well plate (Costar 3631), and optical density (600 nm, OD_600_) and bulk SFGFP fluorescence (485 nm excitation, 520 nm emission) were measured using a Biotek Synergy H1 plate reader. FL and OD measurements were first corrected by subtracting the mean value of the media blank. FL/OD was calculated for each condition and normalized to the mean FL/OD of the control for the specific randomization experiment.

Selected peptides were sequenced using Sanger sequencing with a primer that specifically amplified the peptide plasmid (gcgtgcaatccatcttgttcaatcat). Sequences that were clean (no mixed peaks) and represented full-length peptides (no truncated sequences, insertions, or deletions) were cloned into plasmids using inverse PCR. Amino acid sequences for the peptides that resulted from randomization are listed in Table S3.

### Bulk fluorescence experiments and culturing conditions

Plasmids were transformed into chemically competent *E. coli* MG1655, plated onto LB agar plates containing the appropriate antibiotics, and incubated overnight at 37°C. Individual colonies from each condition were inoculated into 300 μl of LB with the appropriate antibiotics in a 2 ml 96-well block. The block was sealed with a breathe-easier membrane and grown for 17 hours overnight at 37°C while shaking at 100 rpm (Labnet Vortemp 56). Four microliters of the overnight culture were added to 296 μl of MOPS EZ Rich Defined Medium in a new 96-well block and was incubated for 3 hours under the same conditions as above. After incubation, 100 μl of each culture was transferred to a 96-well plate (Costar 3631), and OD_600_ and bulk SFGFP fluorescence (485 nm excitation, 520 nm emission) were measured using a Biotek Synergy H1 plate reader. Each experiment included two sets of controls: a media blank and *E. coli* transformed with control plasmids. FL and OD measurements were first corrected by subtracting the mean value of the media blank. The ratio of the corrected FL and OD (FL/OD) was calculated for each culture. The *E. coli* cultures transformed with control plasmids were then used to correct for autofluorescence by subtracting the average FL/OD from the control cultures from each test culture.

### Minimum inhibitory concentration (MIC) assay

*E. coli* MG1655 cells containing the p_Lux_-peptide constructs were plated onto LB agar plates containing the appropriate antibiotics (50 μg/ml kan for ColE1 plasmids or 25 μg/ml cm for RSF1010 plasmids). Individual colonies were used to inoculate 5 ml of MOPS EZ Rich Defined Medium supplemented with the appropriate antibiotics and cultures were grown for 17 hours at 37°C while shaking at 275 rpm (Shel Lab SSI3). Cell pellets of the overnight cultures were collected by centrifugation of 250 μl at 15,000xg for 1 minute. Cell pellets were resuspended in 1 ml of fresh MOPS medium, and OD_600_ was measured. Cultures were diluted appropriately in MOPS medium to reach a final concentration of 5 x 10^5^ CFU/ml in each well of a 96-well plate (Corning 3370). AHL (Cayman Chemical 143537-62-6) was added to each well at a final concentration of 10 nM for the ColE1 plasmids and 20 nM for the RSF1010 plasmids. A 2-fold serial dilution of the antibiotic was made across the columns of the plate. Cephalothin (Sigma-Aldrich C4520) concentrations ranged from 256 to 0.5 μg/ml. Plates were sealed with a breathe-easier membrane (USA Scientific 9126-2100) followed by a breath-easy membrane (Diversified Biotech, BEM-1) and incubated for 24 hours at 37°C while shaking at 100 rpm (Labnet Vortemp 56). After incubation, the OD_600_ was measured using the Biotek Synergy H1 plate reader. The MIC was determined as a 99% reduction in OD_600_.

### Conjugation assay

The LuxR-p_Lux_-peptide constructs were cloned into the plasmid pRL1342 (RSF1010 origin, Addgene 70691) using restriction enzyme cloning with XbaI and PstI. Recombinant plasmids were transformed into chemically competent *E. coli* WM3064 to create the donor strain. Both donor strain and *E. coli* MG1655 were cultured for 17 hours overnight in LB medium at 37°C while shaking at 275 rpm (Shel Lab SSI3). WM3064 was grown in the presence of 0.3 mM DAP and 25 μg/ml cm. Overnight cultures were subcultured by transferring 60 μl of the culture into 6 ml of fresh LB medium and incubated until the cultures reached an optical density between 0.4 and 0.6. One ml of each culture was collected and centrifuged at 825xg for 5 minutes. Cell pellets were resuspended in 500 μl of LB medium and spun again at 825xg for 5 minutes. Cell pellets were again resuspended in 500 μl of LB medium. Donor and recipient cells were then mixed in a 1:1 ratio (50 μl each) and spotted onto an LB agar plate containing 0.3 mM DAP. After a 1-hour incubation at 37°C, cells were streaked onto an LB agar plate containing 25 μg/ml cm to allow only the successful recipient cells to grow. Upon overnight incubation at 37°C, colony PCR was performed to confirm the presence of the peptides (primer sequences: tggaggccatcaaaccacg, gccattgggatatatcaacggtgg).

### Molecular dynamics simulations

Protein Data Bank (pdb)^53^ files containing needed structures were either created using PyMOL^54^ or obtained from their respective web server. The resolved bacteriophage λ N-protein-NutboxB-RNA complex (PDB ID: 1QFQ)^55^ deposited in the PDB served as the source for the λ N ARM structure. An isolated wild type λ N peptide structure was modeled with SWISS-MODEL^56^, using PDB entry 6GOV^57^ as a template, and exported to a new pdb file for use in subsequent simulations. Using PyMOL’s mutagenesis tool, the λ N peptide was mutated to create the G28_FH35, G28_FH262, and G8 mutants resulting in a new pdb file for each mutant. Because a resolved 3-dimensional structure for the MicF RNA was not available, one was generated using the 3dRNA/DNA web server^58^. The predicted structure with the lowest 3dRNA score (indicative of the best model) was saved for use as MicF’s structure for subsequent steps. The 5’ guanine (G5) and 3’ uracil (U3) of MicF were renamed G and U, respectively, in PyMOL for proper residue recognition by CHARMM-GUI^59–62^. The structures of the wild type λ N peptide and MicF were used to construct a complex for the negative control using ClusPro2.0^63–66^. The model with the most populated cluster, model 0, was chosen, as it represents the most likely λ N-MicF complex. The process was repeated to produce G28_FH35-MicF, G28_FH262-MicF, and G8-MicF models.

CHARMM-GUI’s Solution Builder input generator was utilized to prepare solvated peptide-RNA complex files for MD simulations. Each pdb file was uploaded to the CHARMM-GUI server using the Protein Solution System. All RNA and Protein chains were selected at the Model/Chain selection step to account completely for the desired complexes. Because the peptide contained cysteine residues, a disulfide bond between residues 12 and 19 of G28_FH262 was explicitly added. The system was constrained within a rectangular water box; the size of which was fitted to the protein size and given an edge distance of 10.0 Å. To simulate physiological conditions inside cells, where molecular systems are surrounded by an aqueous environment rather than being confined to a box: K+ and Cl-ions were placed using the Monte-Carlo ion placement method to neutralize the system at a buffer concentration of 0.15 M and the Periodic Boundary Conditions (PBC) were used. The solvated molecular system was simulated using the following physics-based AMBER force fields: FF14SB for Protein, OL15 for DNA, OL3 for RNA, and OPC for Water. The resulting zipped files were downloaded for use in our MD simulations.

AMBER16^67,68^ was used to run the MD simulations of the peptide-RNA complexes. This involved six steps outlined in Santiago and Abrol^69^: 1) Energy minimization of the solvent to minimize its energy and remove protein-solvent steric clashes. 2) Equilibration of the solvent to relax the solvent and allow for artificial air bubbles at the solute-solvent interface to be filled in by the solvent. 3) Energy minimization of the full system to allow the protein atoms to adjust to the newly relaxed solvent. 4) Heating of the full system, which involved the slow heating of the system from 0 K to 310.15 K at 1 atm pressure. For these first four steps, the peptide-RNA complex remained constrained. 5) Equilibration of the full system, wherein the system was relaxed until its density converged. 6) Production run of the full system to simulate potential peptide-RNA binding conformations. One snapshot was taken of the system every 100 picoseconds. A total of 1,000 snapshots were taken, representing 100 nanoseconds of simulation time, a long-enough timespan to test the stability of the peptide-MicF complex. For the λ N peptide, G28_FH35, G28_FH262, and G8 complexes, the simulation was extended for a total of 10,000 snapshots, representing 1 μs of simulated time.

The results of the simulations were visualized with the Visual Molecular Dynamics (VMD)^70^ software program. The completed simulations were analyzed with MMPBSA.py^72^ using the Molecular Mechanics/Poisson-Boltzmann Surface Area (MMPBSA) method^73^ to characterize each complex’s energy profile. To perform the protein-RNA contact analysis, CPPTRAJ^74^, a program for the analysis of MD trajectory data, was used.

## ACKNOWLEDGEMENTS

Research reported in this work was supported by the National Institute of General Medical Sciences of the National Institutes of Health under award number SC2GM136500. The content is solely the responsibility of the authors and does not necessarily represent the official views of the National Institutes of Health.

## References

(1) Beisel, C. L.; Storz, G. Base Pairing Small RNAs and Their Roles in Global Regulatory Networks. FEMS Microbiol Rev 2010, 34 (5), 866–882. 10.1111/j.1574-6976.2010.00241.x.

(2) Storz, G.; Vogel, J.; Wassarman, K. M. Regulation by Small RNAs in Bacteria: Expanding Frontiers. Mol Cell 2011, 43 (6), 880–891. 10.1016/j.molcel.2011.08.022.

(3) Papenfort, K.; Melamed, S. Small RNAs, Large Networks: Posttranscriptional Regulons in Gram-Negative Bacteria. Annual Review of Microbiology 2023, 77 (Volume 77, 2023), 23–43. 10.1146/annurev-micro-041320-025836.

(4) Hör, J.; Matera, G.; Vogel, J.; Gottesman, S.; Storz, G. Trans-Acting Small RNAs and Their Effects on Gene Expression in *Escherichia Coli* and *Salmonella Enterica*. EcoSal Plus 2020, 9 (1), 10.1128/ecosalplus.ESP-0030–2019. 10.1128/ecosalplus.esp-0030-2019.

(5) Mika, F.; Hengge, R. Small Regulatory RNAs in the Control of Motility and Biofilm Formation in *E. Coli* and *Salmonella*. International Journal of Molecular Sciences 2013, 14 (3), 4560–4579. 10.3390/ijms14034560.

(6) Dersch, P.; Khan, M. A.; Mühlen, S.; Görke, B. Roles of Regulatory RNAs for Antibiotic Resistance in Bacteria and Their Potential Value as Novel Drug Targets. Front Microbiol 2017, 8, 803. 10.3389/fmicb.2017.00803.

(7) Henderson, C. A.; Vincent, H. A.; Callaghan, A. J. Reprogramming Gene Expression by Targeting RNA-Based Interactions: A Novel Pipeline Utilizing RNA Array Technology. ACS Synth. Biol. 2021, 10 (8), 1847–1858. 10.1021/acssynbio.0c00603.

(8) Tsai, M. J.; Zambrano, R. A. I.; Susas, J. L.; Silva, L.; Takahashi, M. K. Identifying Antisense Oligonucleotides to Disrupt Small RNA Regulated Antibiotic Resistance via a Cell-Free Transcription–Translation Platform. ACS Synth. Biol. 2023, 12 (8), 2245–2251. 10.1021/acssynbio.3c00245.

(9) Good, L.; Awasthi, S. K.; Dryselius, R.; Larsson, O.; Nielsen, P. E. Bactericidal Antisense Effects of Peptide–PNA Conjugates. Nat Biotechnol 2001, 19 (4), 360–364. 10.1038/86753.

(10) Goltermann, L.; Yavari, N.; Zhang, M.; Ghosal, A.; Nielsen, P. E. PNA Length Restriction of Antibacterial Activity of Peptide-PNA Conjugates in Escherichia Coli Through Effects of the Inner Membrane. Frontiers in Microbiology 2019, 10. 10.3389/fmicb.2019.01032.

(11) Gottesman, M. E.; Adhya, S.; Das, A. Transcription Antitermination by Bacteriophage Lambda *N* Gene Product. Journal of Molecular Biology 1980, 140 (1), 57–75. 10.1016/0022-2836(80)90356-3.

(12) Frankel, A. D.; Young, J. A. T. HIV-1: Fifteen Proteins and an RNA. Annu. Rev. Biochem. 1998, 67 (1), 1–25. 10.1146/annurev.biochem.67.1.1.

(13) Tan, R.; Chen, L.; Buettner, J. A.; Hudson, D.; Frankel, A. D. RNA Recognition by an Isolated α Helix. Cell 1993, 73 (5), 1031–1040. 10.1016/0092-8674(93)90280-4.

(14) Tan, R.; Frankel, A. D. Structural Variety of Arginine-Rich RNA-Binding Peptides. Proceedings of the National Academy of Sciences 1995, 92 (12), 5282–5286. 10.1073/pnas.92.12.5282.

(15) Austin, R. J.; Xia, T.; Ren, J.; Takahashi, T. T.; Roberts, R. W. Designed Arginine-Rich RNA-Binding Peptides with Picomolar Affinity. J. Am. Chem. Soc. 2002, 124 (37), 10966–10967. 10.1021/ja026610b.

(16) Harada, K.; Martin, S. S.; Tan, R.; Frankel, A. D. Molding a Peptide into an RNA Site by in Vivo Peptide Evolution. Proceedings of the National Academy of Sciences 1997, 94 (22), 11887–11892. 10.1073/pnas.94.22.11887.

(17) Harada, K.; Martin, S. S.; Frankel, A. D. Selection of RNA-Binding Peptides in Vivo. Nature 1996, 380 (6570), 175–179. 10.1038/380175a0.

(18) Mizuno, T.; Chou, M. Y.; Inouye, M. A Unique Mechanism Regulating Gene Expression: Translational Inhibition by a Complementary RNA Transcript (micRNA). Proc Natl Acad Sci 1984, 81 (7), 1966–1970. 10.1073/pnas.81.7.1966.

(19) Andersen, J.; Forst, S. A.; Zhao, K.; Inouye, M.; Delihas, N. The Function of *micF* RNA. *micF* RNA Is a Major Factor in the Thermal Regulation of OmpF Protein in *Escherichia Coli*. J Biol Chem 1989, 264 (30), 17961–17970. 10.1016/S0021-9258(19)84666-5.

(20) Hächler, H.; Cohen, S. P.; Levy, S. B. *marA*, a Regulated Locus Which Controls Expression of Chromosomal Multiple Antibiotic Resistance in *Escherichia Coli*. J Bacteriol 1991, 173 (17), 5532–5538. 10.1128/jb.173.17.5532-5538.1991.

(21) Oh, J. T.; Cajal, Y.; Skowronska, E. M.; Belkin, S.; Chen, J.; Van Dyk, T. K.; Sasser, M.; Jain, M. K. Cationic Peptide Antimicrobials Induce Selective Transcription of *micF* and *osmY* in *Escherichia Coli*. Biochim Biophys Acta 2000, 1463 (1), 43–54. 10.1016/s0005-2736(99)00177-7.

(22) Chou, J. H.; Greenberg, J. T.; Demple, B. Posttranscriptional Repression of *Escherichia Coli* OmpF Protein in Response to Redox Stress: Positive Control of the *micF* Antisense RNA by the *soxRS* Locus. J Bacteriol 1993, 175 (4), 1026– 1031. 10.1128/jb.175.4.1026-1031.1993.

(23) Kim, T.; Bak, G.; Lee, J.; Kim, K. Systematic Analysis of the Role of Bacterial Hfq-Interacting sRNAs in the Response to Antibiotics. Journal of Antimicrobial Chemotherapy 2015, 70 (6), 1659–1668. 10.1093/jac/dkv042.

(24) Delihas, N.; Forst, S. *MicF*: An Antisense RNA Gene Involved in Response of *Escherichia Coli* to Global Stress Factors. J Mol Biol 2001, 313 (1), 1–12. 10.1006/jmbi.2001.5029.

(25) Nikaido, H. Prevention of Drug Access to Bacterial Targets: Permeability Barriers and Active Efflux. Science 1994, 264 (5157), 382–388. 10.1126/science.8153625.

(26) Choi, U.; Lee, C.-R. Distinct Roles of Outer Membrane Porins in Antibiotic Resistance and Membrane Integrity in *Escherichia Coli*. Frontiers in Microbiology 2019, 10. 10.3389/fmicb.2019.00953.

(27) Corcoran, C. P.; Podkaminski, D.; Papenfort, K.; Urban, J. H.; Hinton, J. C. D.; Vogel, J. Superfolder GFP Reporters Validate Diverse New mRNA Targets of the Classic Porin Regulator, MicF RNA. Mol Microbiol 2012, 84 (3), 428–445. 10.1111/j.1365-2958.2012.08031.x.

(28) Registry of Standard Biological Parts - Promoters/Catalog/Anderson. https://parts.igem.org/Promoters/Catalog/Anderson (accessed 2025-05-06).

(29) Weiss, M. A.; Narayana, N. RNA Recognition by Arginine-Rich Peptide Motifs. Biopolymers 1998, 48 (2–3), 167–180. 10.1002/(SICI)1097-0282(1998)48:2%3C167::AID-BIP6%3E3.0.CO;2-8.

(30) Cocozaki, A. I.; Ghattas, I. R.; Smith, C. A. The RNA-Binding Domain of Bacteriophage P22 N Protein Is Highly Mutable, and a Single Mutation Relaxes Specificity toward λ. Journal of Bacteriology 2008, 190 (23), 7699–7708. 10.1128/jb.00997-08.

(31) Chen, L.; Frankel, A. D. An RNA-Binding Peptide from Bovine Immunodeficiency Virus Tat Protein Recognizes an Unusual RNA Structure. Biochemistry 1994, 33 (9), 2708–2715. 10.1021/bi00175a046.

(32) Canton, B.; Labno, A.; Endy, D. Refinement and Standardization of Synthetic Biological Parts and Devices. Nat Biotechnol 2008, 26 (7), 787–793. 10.1038/nbt1413.

(33) Drayton, M.; Kizhakkedathu, J. N.; Straus, S. K. Towards Robust Delivery of Antimicrobial Peptides to Combat Bacterial Resistance. Molecules 2020, 25 (13), 3048. 10.3390/molecules25133048.

(34) Citorik, R. J.; Mimee, M.; Lu, T. K. Sequence-Specific Antimicrobials Using Efficiently Delivered RNA-Guided Nucleases. Nat Biotechnol 2014, 32 (11), 1141–1145. 10.1038/nbt.3011.

(35) Lam, K. N.; Spanogiannopoulos, P.; Soto-Perez, P.; Alexander, M.; Nalley, M. J.; Bisanz, J. E.; Nayak, R. R.; Weakley, A. M.; Yu, F. B.; Turnbaugh, P. J. Phage-Delivered CRISPR-Cas9 for Strain-Specific Depletion and Genomic Deletions in the Gut Microbiome. Cell Reports 2021, 37 (5), 109930. 10.1016/j.celrep.2021.109930.

(36) Huan, Y. W.; Torraca, V.; Brown, R.; Fa-arun, J.; Miles, S. L.; Oyarzún, D. A.; Mostowy, S.; Wang, B. P1 Bacteriophage-Enabled Delivery of CRISPR-Cas9 Antimicrobial Activity Against Shigella Flexneri. ACS Synth. Biol. 2023, 12 (3), 709–721. 10.1021/acssynbio.2c00465.

(37) Sheng, H.; Wu, S.; Xue, Y.; Zhao, W.; Caplan, A. B.; Hovde, C. J.; Minnich, S. A. Engineering Conjugative CRISPR-Cas9 Systems for the Targeted Control of Enteric Pathogens and Antibiotic Resistance. PLoS One 2023, 18 (9), e0291520. 10.1371/journal.pone.0291520.

(38) Derollez, E.; Lesterlin, C.; Bigot, S. Design, Potential and Limitations of Conjugation-based Antibacterial Strategies. Microb Biotechnol 2024, 17 (11), e70050. 10.1111/1751-7915.70050.

(39) Schu, D. J.; Zhang, A.; Gottesman, S.; Storz, G. Alternative Hfq-sRNA Interaction Modes Dictate Alternative mRNA Recognition. The EMBO Journal 2015, 34 (20), 2557–2573. 10.15252/embj.201591569.

(40) Urban, J. H.; Vogel, J. Translational Control and Target Recognition by *Escherichia Coli* Small RNAs *in Vivo*. Nucleic Acids Res 2007, 35 (3), 1018–1037. 10.1093/nar/gkl1040.

(41) Stibelman, A. Y.; Sariles, A. Y.; Takahashi, M. K. The Small RNA MicF Represses ObgE and SeqA in *Escherichia Coli*. Microorganisms 2024, 12 (12), 2397. 10.3390/microorganisms12122397.

(42) Papenfort, K.; Bouvier, M.; Mika, F.; Sharma, C. M.; Vogel, J. Evidence for an Autonomous 5′ Target Recognition Domain in an Hfq-Associated Small RNA. Proc Natl Acad Sci 2010, 107 (47), 20435–20440. 10.1073/pnas.1009784107.

(43) Fouts, D. E.; Celander, D. W. Improved Method for Selecting RNA-Binding Activities in Vivo. Nucleic Acids Res 1996, 24 (8), 1582–1584. 10.1093/nar/24.8.1582.

(44) Wilhelm, J. E.; Vale, R. D. A One-Hybrid System for Detecting RNA–Protein Interactions. Genes to Cells 1996, 1 (3), 317–323. 10.1046/j.1365-2443.1996.25026.x.

(45) SenGupta, D. J.; Zhang, B.; Kraemer, B.; Pochart, P.; Fields, S.; Wickens, M. A Three-Hybrid System to Detect RNA-Protein Interactions *in Vivo*. Proceedings of the National Academy of Sciences 1996, 93 (16), 8496–8501. 10.1073/pnas.93.16.8496.

(46) Laird-Offringa, I. A.; Belasco, J. G. Analysis of RNA-Binding Proteins by *in Vitro* Genetic Selection: Identification of an Amino Acid Residue Important for Locking U1A onto Its RNA Target. Proceedings of the National Academy of Sciences 1995, 92 (25), 11859–11863. 10.1073/pnas.92.25.11859.

(47) Parker, A.; Gottesman, S. Small RNA Regulation of TolC, the Outer Membrane Component of Bacterial Multidrug Transporters. Journal of Bacteriology 2016, 198 (7), 1101–1113. 10.1128/jb.00971-15.

(48) Liu, J.; Cropp, T. A. A Method for Multi-Codon Scanning Mutagenesis of Proteins Based on Asymmetric Transposons. Protein Eng Des Sel 2012, 25 (2), 67–72. 10.1093/protein/gzr060.

(49) Stacey, S. B.; Sechkar, K.; Corrao, M.; Steel, H.; Papachristodoulou, A. Quantitative Engineering and Investigation of Synthetic Sponge RNAs in *E*. Coli. bioRxiv May 20, 2026, p 2026.05.19.726096. 10.64898/2026.05.19.726096.

(50) Gadar, K.; McCarthy, R. R. Using next Generation Antimicrobials to Target the Mechanisms of Infection. npj Antimicrob Resist 2023, 1 (1), 1–14. 10.1038/s44259-023-00011-6.

(51) Dehbanipour, R.; Ghalavand, Z. Anti-Virulence Therapeutic Strategies against Bacterial Infections: Recent Advances. Germs 2022, 12 (2), 262–275. 10.18683/germs.2022.1328.

(52) Ogawara, H. Possible Drugs for the Treatment of Bacterial Infections in the Future: Anti-Virulence Drugs. J Antibiot 2021, 74 (1), 24–41. 10.1038/s41429-020-0344-z.

(53) Berman, H. M. The Protein Data Bank. Nucleic Acids Research 2000, 28 (1), 235–242. 10.1093/nar/28.1.235.

(54) The PyMOL Molecular Graphics System, Version 2.6 Schrodinger, LLC.

(55) Schärpf, M.; Sticht, H.; Schweimer, K.; Boehm, M.; Hoffmann, S.; Rösch, P. Antitermination in Bacteriophage λ. European Journal of Biochemistry 2000, 267 (8), 2397–2408. 10.1046/j.1432-1327.2000.01251.x.

(56) Waterhouse, A.; Bertoni, M.; Bienert, S.; Studer, G.; Tauriello, G.; Gumienny, R.; Heer, F. T.; Beer, T. A. P. de; Rempfer, C.; Bordoli, L.; Lepore, R.; Schwedeo, T. SWISS-MODEL: Homology Modelling of Protein Structures and Complexes. Nucleic Acids Research 2018, 46 (W1), W296–W303. 10.1093/nar/gky427.

(57) Krupp, F.; Said, N.; Huang, Y.-H.; Loll, B.; Bürger, J.; Mielke, T.; Spahn, C. M. T.; Wahl, M. C. Structural Basis for the Action of an All-Purpose Transcription Anti-Termination Factor. Molecular Cell 2019, 74 (1), 143–157.e5. 10.1016/j.molcel.2019.01.016.

(58) Zhang, Y.; Wang, J.; Xiao, Y. 3dRNA: 3D Structure Prediction from Linear to Circular RNAs. Journal of Molecular Biology 2022, 434 (11), 167452. 10.1016/j.jmb.2022.167452.

(59) Jo, S.; Kim, T.; Iyer, V. G.; Im, W. CHARMM-GUI: A Web-Based Graphical User Interface for CHARMM. Journal of Computational Chemistry 2008, 29 (11), 1859–1865. 10.1002/jcc.20945.

(60) Brooks, B. R.; Brooks, C. L.; Mackerell, A. D.; Nilsson, L.; Petrella, R. J.; Roux, B.; Won, Y.; Archontis, G.; Bartels, C.; Boresch, S.; Caflisch, A.; Caves, L.; Cui, Q.; Dinner, A. R.; Feig, M.; Fischer, S.; Gao, J.; Hodoscek, M.; Im, W.; Kuczera, K.; Lazaridis, T.; Ma, J.; Ovchinnikov, V.; Paci, E.; Pastor, R. W.; Post, C. B.; Pu, J. Z.; Schaefer, M.; Tidor, B.; Venable, R. M.; Woodcock, H. L.; Wu, X.; Yang, W.; York, D. M.; Karplus, M. CHARMM: The Biomolecular Simulation Program. Journal of Computational Chemistry 2009, 30 (10), 1545–1614. 10.1002/jcc.21287.

(61) Lee, J.; Cheng, X.; Swails, J. M.; Yeom, M. S.; Eastman, P. K.; Lemkul, J. A.; Wei, S.; Buckner, J.; Jeong, J. C.; Qi, Y.; Jo, S.; Pande, V. S.; Case, D. A.; Brooks, C. L.; MacKerell, A. D.; Klauda, J. B.; Im, W. CHARMM-GUI Input Generator for NAMD, GROMACS, AMBER, OpenMM, and CHARMM/OpenMM Simulations Using the CHARMM36 Additive Force Field. Journal of Chemical Theory and Computation 2016, 12 (1), 405–413. 10.1021/acs.jctc.5b00935.

(62) Lee, J.; Hitzenberger, M.; Rieger, M.; Kern, N. R.; Zacharias, M.; Im, W. CHARMM-GUI Supports the Amber Force Fields. The Journal of Chemical Physics 2020, 153 (3), 035103. 10.1063/5.0012280.

(63) Kozakov, D.; Beglov, D.; Bohnuud, T.; Mottarella, S. E.; Xia, B.; Hall, D. R.; Vajda, S. How Good Is Automated Protein Docking? Proteins: Structure, Function, and Bioinformatics 2013, 81 (12), 2159–2166. 10.1002/prot.24403.

(64) Vajda, S.; Yueh, C.; Beglov, D.; Bohnuud, T.; Mottarella, S. E.; Xia, B.; Hall, D. R.; Kozakov, D. New Additions to the ClusPro Server Motivated by CAPRI. Proteins: Structure, Function, and Bioinformatics 2017, 85 (3), 435–444. 10.1002/prot.25219.

(65) Kozakov, D.; Hall, D. R.; Xia, B.; Porter, K. A.; Padhorny, D.; Yueh, C.; Beglov, D.; Vajda, S. The ClusPro Web Server for Protein–Protein Docking. Nature Protocols 2017, 12 (2), 255–278. 10.1038/nprot.2016.169.

(66) Desta, I. T.; Porter, K. A.; Xia, B.; Kozakov, D.; Vajda, S. Performance and Its Limits in Rigid Body Protein-Protein Docking. Structure 2020, 28 (9), 1071–1081.e3. 10.1016/j.str.2020.06.006.

(67) Case, D. A.; Cheatham, T. E.; Darden, T.; Gohlke, H.; Luo, R.; Merz, K. M.; Onufriev, A.; Simmerling, C.; Wang, B.; Woods, R. J. The Amber Biomolecular Simulation Programs. Journal of Computational Chemistry 2005, 26 (16), 1668–1688. 10.1002/jcc.20290.

(68) Salomon-Ferrer, R.; Case, D. A.; Walker, R. C. An Overview of the Amber Biomolecular Simulation Package. Wiley Interdisciplinary Reviews: Computational Molecular Science 2013, 3 (2), 198–210. 10.1002/wcms.1121.

(69) Santiago, L.; Abrol, R. Understanding G Protein Selectivity of Muscarinic Acetylcholine Receptors Using Computational Methods. International Journal of Molecular Sciences 2019, 20 (21), 5290. 10.3390/ijms20215290.

(70) Humphrey, W.; Dalke, A.; Schulten, K. VMD: Visual Molecular Dynamics. Journal of Molecular Graphics 1996, 14 (1), 33–38. 10.1016/0263-7855(96)00018-5.

(71) Adasme, M. F.; Linnemann, K. L.; Bolz, S. N.; Kaiser, F.; Salentin, S.; Haupt, V. J.; Schroeder, M. PLIP 2021: Expanding the Scope of the Protein–Ligand Interaction Profiler to DNA and RNA. Nucleic Acids Research 2021, 49 (W1), W530–W534. 10.1093/nar/gkab294.

(72) Miller, B. R.; McGee, T. D.; Swails, J. M.; Homeyer, N.; Gohlke, H.; Roitberg, A. E. MMPBSA.Py : An Efficient Program for End-State Free Energy Calculations. Journal of Chemical Theory and Computation 2012, 8 (9), 3314–3321. 10.1021/ct300418h.

(73) Wang, C.; Greene, D.; Xiao, L.; Qi, R.; Luo, R. Recent Developments and Applications of the MMPBSA Method. Frontiers in Molecular Biosciences 2018, 4. 10.3389/fmolb.2017.00087.

(74) Roe, D. R.; Cheatham, T. E. PTRAJ and CPPTRAJ: Software for Processing and Analysis of Molecular Dynamics Trajectory Data. Journal of Chemical Theory and Computation 2013, 9 (7), 3084–3095. 10.1021/ct400341p.

