## Supporting Information for "Disruption of sRNA Function Using Synthetic Arginine Rich Motif Peptides"

Table of contents:


| | **Description** | **Page** |
|---|---|---|
| Table S1 | Important DNA sequences | 2-4 |
| Table S2 | Primer sequences for randomization cloning | 4 |
| Table S3 | Amino acid sequences for peptides from randomizations | 5 |
| Table S4 | Plasmids used in this study | 6-7 |
| Figure S1 | Additional randomization results for BIV-Tat, P22-N, and HIV-REV | 8 |
| Figure S2 | Randomization of peptide λ N G24 | 9 |
| Figure S3 | Randomization of peptide λ N G31 | 10 |
| Figure S4 | Randomization of peptide λ N G32 | 11 |
| Figure S5 | PCR to confirm conjugation of peptide carrying plasmid | 12 |
| Figure S6 | Free energy analysis of peptide-MicF complexes at 10,000 frames | 12 |
| Table S5 | Electrostatic energy contribution per peptide residue in λ N-MicF MD simulation from MMPBSA analysis | 13 |
| Table S6 | Electrostatic energy contribution per peptide residue in G8-MicF MD simulation from MMPBSA analysis | 14 |
| Table S7 | Electrostatic energy contribution per peptide residue in G28_FH35-MicF MD simulation from MMPBSA analysis | 15 |
| Table S8 | Electrostatic energy contribution per peptide residue in G28_FH262-MicF MD simulation from MMPBSA analysis | 16 |
| Table S9 | AA-nucleotide Contact analysis for λ N-MicF MD simulation | 17-18 |
| Table S10 | AA-nucleotide Contact analysis for G8-MicF MD simulation | 19 |
| Table S11 | AA-nucleotide Contact analysis for G28_FH35-MicF MD simulation | 20 |
| Table S12 | AA-nucleotide Contact analysis for G28_FH262-MicF MD simulation | 21-22 |

**Table S1**: Important DNA sequences

| Name | Sequence |
|---|---|
| pLux promoter | acctgtaggatcgtacaggtttacgcaagaaaatggtttgttatagtcgaataaa |
| LuxR sequence | atgaaaaacataaatgccgacgacacatacagaataattaataaaattaaagcttgtagaagcaataatgatattaatcaatgcttatctgatatgactaaaatggtacattgtgaatattatttactcgcgatcatttatcctcattctatggttaaatctgatatttcaatcctagataattaccctaaaaaatggaggcaatattatgatgacgctaatttaataaaatatgatcctatagtagattattctaactccaatcattcaccaattaattggaatatatttgaaaacaatgctgtaaataaaaaatctccaaatgtaattaaagaagcgaaaacatcaggtcttatcactgggtttagtttcccattcatacggctaacaatggcttcggaatgcttagttttgcacattcagaaaaagacaactatatagatagtttatttttacatgcgtgtatgaacataccattaattgttccttctctagttgataattatcgaaaaataaatatagcaaataataaatcaaacaacgatttaaccaaaagagaaaaagaatgtttagcgtgggcatgcgaaggaaaaagctcttgggatatttcaaaaatattaggttgcagtgagcgtactgtcactttccatttaaccaatgcgcaaatgaaactcaatacaacaaaccgctgccaaagtatttctaaagcaattttaacaggagcaattgattgcccatactttaaaaattaa |
| J23118 promoter | ttgacggctagctcagtcctaggtattgtgctagc |
| ompF-sfgfp fusion | agacacataaagacaccaaactctcatcaatagttccgtaaatttttattgacagaacttattgacggcagtggcaggtgtcataaaaaaaaccatgagggtaataaataatgatgaagcgcaatattctggcagtgatcgtccctgctagcaaaggagaagaacttttcactggagttgtcccaattcttgttgaattagatggtgatgttaatgggcacaaattttctgtccgtggagagggtgaaggtgatgctacaaacggaaaactcacccttaaatttatttgcactactggaaaactacctgttccgtggccaacacttgtcactactctgacctatggtgttcaatgcttttcccgttatccggatcacatgaaacggcatgactttttcaagagtgccatgcccgaaggttatgtacaggaacgcactatatctttcaaagatgacgggacctacaagacgcgtgctgaagtcaagtttgaaggtgatacccttgttaatcgtatcgagttaaagggtattgattttaaagaagatggaaacattcttggacacaaactcgagtacaactttaactcacacaatgtatacatcacggcagacaaacaaaagaatggaatcaaagctaacttcaaaattcgccacaacgttgaagatggttccgttcaactagcagaccattatcaacaaaatactccaattggcgatggccctgtccttttaccagacaaccattacctgtcgacacaatctgtcctttcgaaagatcccaacgaaaagcgtgaccacatggtccttcttgagtttgtaactgctgctgggattacacatggcatggatgagctctacaaataa |
| MicF | gctatcatcattaactttatttattaccgtcattcatttctgaatgtctgtttacccctatttcaaccggatgcctcgcattcggtttttttt |
| T1 terminator | gcatcaaataaaacgaaaggctcagtcgaaagactgggcctttcgttttatctgttgtttgtcggtgaacgctctcctgagtaggacaaatccgccgccctaga |
| SgrS scaffold (SS) | gctatcatcattatattggtgtaaaatcacccgccagcagattatacctgctggtttttttt |
| MicF(1-13) SS | gctatcatcattatattggtgtaaaatcacccgccagcagattatacctgctggtttttttt |
| RybB | gccactgcttttctttgatgtccccattttgtggagcccatcaaccccgccatttcggttcaaggttgatgggttttttgt |
| Lrp-sfgfp fusion | ggaagaaaaaaaacagtattcttatatgcgcataaccatgcatgtaaatataccatgtttaccgtgctagtgaaatctacgtatggcgtggacagacgccattcgt |

| | |
|---|---|
| | gatgtcgatagctgccacaaggcaacggtcttctcaccgtagacccaggc attgcgcgccgtgaatcttcatgatttcggtctatcgtgacgggtagcgactct gaacagtgatgtttcagggtcagacaggagtagggaaggaatacagaga gacaataataatggtagatagcaagaagcgccctggcaaagatctcgac cgtatcgatcgtaacattcttagcaaaggagaagaacttttcactggagttgt cccaattcttgttgaattagatggtgatgttaatgggcacaaattttctgtccgt ggagagggtgaaggtgatgctacaaacggaaaactcacccttaaatttatt tgcactactggaaaactacctgttccgtggccaacacttgtcactactctgac ctatggtgttcaatgcttttcccgttatccggatcacatgaaacggcatgactt ttcaagagtgccatgcccgaaggttatgtacaggaacgcactatatctttca aagatgacgggacctacaagacgcgtgctgaagtcaagtttgaaggtgat acccttgttaatcgtatcgagttaaagggtattgattttaaagaagatggaaa cattcttggacacaaactcgagtacaactttaactcacacaatgtatacatc acggcagacaaacaaaagaatggaatcaaagctaacttcaaaattcgcc acaacgttgaagatggttccgttcaactagcagaccattatcaacaaaata ctccaattggcgatggccctgtccttttaccagacaaccattacctgtcgaca caatctgtcctttcgaaagatcccaacgaaaagcgtgaccacatggtccttc ttgagtttgtaactgctgctgggattacacatggcatggatgagctctacaaa taa |
| obgE-sfgfp fusion | gaattcattaaagaggagaaaggtaccatggactacaaagaccatgacg gtgattataaagatcatgatatcgactacaaagatgacgacgataaaacc atgattacggattcactggccgtcgttttacaacgtcgtgactgggaaaacc ctggcgttacccaacttaatcgccttgcagcacatccccctttcgccagctgg cgtaatagcgaagaggcccgcaccgatcgcccttcccaacagttgcgca gcctgaatggcgaatggatgcattaagcatacaacggtggtatcgcaacc ccgcgcaggcgaatgatttacggagaataaaatgaagtttgttgatgaagc atcgattctggtcgttagcaaaggagaagaacttttcactggagttgtcccaa ttcttgttgaattagatggtgatgttaatgggcacaaattttctgtccgtggaga gggtgaaggtgatgctacaaacggaaaactcacccttaaatttatttgcact actggaaaactacctgttccgtggccaacacttgtcactactctgacctatgg tgttcaatgcttttcccgttatccggatcacatgaaacggcatgactttttcaag agtgccatgcccgaaggttatgtacaggaacgcactatatctttcaaagatg acgggacctacaagacgcgtgctgaagtcaagtttgaaggtgatacccttg ttaatcgtatcgagttaaagggtattgattttaaagaagatggaaacattcttg gacacaaactcgagtacaactttaactcacacaatgtatacatcacggca gacaaacaaaagaatggaatcaaagctaacttcaaaattcgccacaacg ttgaagatggttccgttcaactagcagaccattatcaacaaaatactccaatt ggcgatggccctgtccttttaccagacaaccattacctgtcgacacaatctgt cctttcgaaagatcccaacgaaaagcgtgaccacatggtccttcttgagttt gtaactgctgctgggattacacatggcatggatgagctctacaaataa |
| λ N | atggacgcgcagacacgtcggcgcgaacggagagccgaaaagcaag cacaatggaaagctgcaaac |
| BIV-Tat | agcggtcctcgtccgcgtggaacaagagggaaaggtcggagaattcgtc gc |
| P22-N | aacgccaagacacgtcgccatgaacgtcgtcggaaacttgccatcgaac gc |
| HIV-REV | accagacaagctcggcgtaacagaagacgcagatggcgggagcgtca acgc |
| J23118 – RBS λ N – T1 | ttgacggctagctcagtcctaggtattgtgctagcgaattcattaaagagga gaaaggtaccatgatggacgcgcagacacgtcggcgcgaacggagag |

| | |
|---|---|
| | ccgaaaagcaagcacaatggaaagctgcaaactaagtacgcgtgctagaggcatcaaataaaacgaaaggctcagtcgaaagactgggcctttcgttttatctgttgtttgtcggtgaacgctctcctgagtaggacaaatccgccgccctaga |

**Table S2.** Primer sequences for randomization cloning.

| Randomization | Fwd Primer Sequence (5'-3') | Rev Primer Sequence (5'-3') |
|---|---|---|
| BIV-Tat Region A | NNNNNNACGCGGACGAGGACCGCTCATGGTACCTTTCTCCTCTTTAATGAATTCG | AGAGGGAAAGGTCGGAGAATTCGTCGCTAAGTACGCGTGCTAGAGGCAT |
| BIV-Tat Region B | TGTTCCACGCGGACGAGGACCGCTCATGGTACCTTTCTCCTCTTTAATGAATTC | AGANNNAAAGGTCGGAGAATTCGTCGCTAAGTACGCGTGCTAGAGGCAT |
| P22N Region C | CGTCGGAAACTTGCCATCGAACGCTAAGTACGCGTGCTAGAGGCAT | ACGTTCATGGCGACGNNNNNNNNNNNNCATGGTACCTTTCTCCTCTTTAATGAATTCG |
| P22N Region D | ACGTTCATGGCGACGTGTCTTGGCGTTCATGGTACCTTTCTCCTCTTTAATGAATTCG | CGTCGGNNNNNNNNNNNNNNNCGCTAAGTACGCGTGCTAGAGGCAT |
| HIV-REV Region E | AGAAGACGCAGATGGCGGGAGCGTCAACGCTAAGTACGCGTGCTAGAGGCAT | GTTACGCCGNNNNNNTCTNNNCATGGTACCTTTCTCCTCTTTAATGAATTCG |
| λ N Region F | GGAGANNNNNNNNNNNNGCACAATGGAAAGCTGCAAACTAAGTACGCGTGCTAGAGGCAT | GTTCGCGCCGACGTGTCTGCGCGTCCATCATGGTACCTTTCTCCTCTTTAATGAATTCG |
| λ N Region G | GGAGAGCCGAAAAGCAAGCACAATGGAAAGCTGCAAACTAAGTACGCGTGCTAGAGGCAT | GTTCGCGCCGACGNNNNNNNNNNNNCATCATGGTACCTTTCTCCTCTTTAATGAATTCG |
| λ N G24 Region FH | GGAGANNNNNNNNNNNNNNNNNNNNNNNNGCTGCAAACTAAGTACGCGTGCTAGAGGCAT | GTTCGCGCCGACGGATTGCGGCTATCATCATGGTACCTTTCTCCTCTTTAATGAATTCG |
| λ N G28 Region FH | GGAGANNNNNNNNNNNNNNNNNNNNNNNNGCTGCAAACTAAGTACGCGTGCTAGAGGCAT | GTTCGCGCCGACGGTGTTTTGCTATCATCATGGTACCTTTCTCCTCTTTAATGAATTCG |
| λ N G31 Region FH | GGAGANNNNNNNNNNNNNNNNNNNNNNNNGCTGCAAACTAAGTACGCGTGCTAGAGGCAT | GTTCGCGCCGACGGTACTTTGCTATCATCATGGTACCTTTCTCCTCTTT |
| λ N G32 Region FH | GGAGANNNNNNNNNNNNNNNNNNNNNNNNGCTGCAAACTAAGTACGCGTGCTAGAGGCAT | GTTCGCGCCGACGCATACGGGCTATCATCATGGTACCTTTCTCCTCTTT |

**Table S3**. Amino acid sequences for peptides from randomizations.

| | | Region G | | | | | | | | | | Region F | | | | Region H | | | | | | |
|---|---|---|---|---|---|---|---|---|---|---|---|---|---|---|---|---|---|---|---|---|---|---|
| | 1 | 2 | 3 | 4 | 5 | 6 | 7 | 8 | 9 | 10 | 11 | 12 | 13 | 14 | 15 | 16 | 17 | 18 | 19 | 20 | 21 | 22 |
| λ N | M | D | A | Q | T | R | R | R | E | R | R | A | E | K | Q | A | Q | W | K | A | A | N |
| G6 | M | I | A | L | C | R | R | R | E | R | R | A | E | K | Q | A | Q | W | K | A | A | N |
| G8 | M | T | A | I | N | R | R | R | E | R | R | A | E | K | Q | A | Q | W | K | A | A | N |
| G13 | M | M | A | V | K | R | R | R | E | R | R | A | E | K | Q | A | Q | W | K | A | A | N |
| G24 | M | I | A | A | I | R | R | R | E | R | R | A | E | K | Q | A | Q | W | K | A | A | N |
| G28 | M | I | A | K | H | R | R | R | E | R | R | A | E | K | Q | A | Q | W | K | A | A | N |
| G31 | M | I | A | K | Y | R | R | R | E | R | R | A | E | K | Q | A | Q | W | K | A | A | N |
| G32 | M | I | A | R | M | R | R | R | E | R | R | A | E | K | Q | A | Q | W | K | A | A | N |
| G24-FH57 | M | I | A | A | I | R | R | R | E | R | R | R | K | F | S | R | Y | P | L | A | A | N |
| G28-FH5 | M | I | A | K | H | R | R | R | E | R | R | R | C | Q | L | A | V | F | I | A | A | N |
| G28-FH35 | M | I | A | K | H | R | R | R | E | R | R | I | L | F | Y | H | I | R | D | A | A | N |
| G28-FH101 | M | I | A | K | H | R | R | R | E | R | R | Y | F | N | C | L | Y | S | C | A | A | N |
| G28-FH108 | M | I | A | K | H | R | R | R | E | R | R | I | Y | Y | F | S | N | G | A | A | A | N |
| G28-FH109 | M | I | A | K | H | R | R | R | E | R | R | S | Y | G | V | C | V | A | L | A | A | N |
| G28-FH117 | M | I | A | K | H | R | R | R | E | R | R | S | L | F | I | S | R | E | E | A | A | N |
| G28-FH133 | M | I | A | K | H | R | R | R | E | R | R | T | P | L | S | I | L | G | W | A | A | N |
| G28-FH262 | M | I | A | K | H | R | R | R | E | R | R | C | L | L | N | S | L | L | C | A | A | N |
| G31-FH10 | M | I | A | K | Y | R | R | R | E | R | R | F | D | L | R | A | G | I | E | A | A | N |
| G31-FH20 | M | I | A | K | Y | R | R | R | E | R | R | S | L | R | P | S | R | S | C | A | A | N |
| G31-FH28 | M | I | A | K | Y | R | R | R | E | R | R | N | L | L | V | R | L | W | L | A | A | N |
| G31-FH40 | M | I | A | K | Y | R | R | R | E | R | R | S | K | G | S | P | S | E | D | A | A | N |
| G31-FH47 | M | I | A | K | Y | R | R | R | E | R | R | F | C | S | V | A | V | F | G | A | A | N |
| G31-FH53 | M | I | A | K | Y | R | R | R | E | R | R | C | N | C | Y | T | Y | S | A | A | A | N |
| G31-FH81 | M | I | A | K | Y | R | R | R | E | R | R | F | T | V | W | R | C | D | S | A | A | N |
| G31-FH144 | M | I | A | K | Y | R | R | R | E | R | R | S | T | F | V | P | I | R | R | A | A | N |
| G32-FH8 | M | I | A | R | M | R | R | R | E | R | R | C | W | I | N | T | S | R | G | A | A | N |
| G32-FH11 | M | I | A | R | M | R | R | R | E | R | R | Y | N | K | V | L | L | R | M | A | A | N |
| G32-FH112 | M | I | A | R | M | R | R | R | E | R | R | L | F | I | S | W | S | P | R | A | A | N |
| G32-FH152 | M | I | A | R | M | R | R | R | E | R | R | N | I | N | I | C | A | K | V | A | A | N |
| G32-FH164 | M | I | A | R | M | R | R | R | E | R | R | T | C | M | L | N | G | V | L | A | A | N |

**Table S4**. Plasmids used in this study.

| Plasmid # | Description | Origin/Resistance | Figure |
|---|---|---|---|
| MKT221 | J23118 – T1 (control) | pSC101/CmR | 1, 2, 5, S2, S3, S4 |
| MKT176 | J23118 – T1 (control) | p15A/AmpR | 1, 2, 5, S2, S3, S4 |
| MKT178 | J23118 – T1 (control) | ColE1/KanR | 1, 2, 5, S2, S3, S4 |
| MKT172 | J23118 – ompF-sfgfp – T1 | pSC101/CmR | 1, 2, 5, S1, S2, S3, S4 |
| MKT173 | J23118 – MicF – T1 | p15A/AmpR | 1, 2, 5, S1, S2, S3, S4 |
| MKT174 | J23118 – antiMicF(33-1) – T1 | ColE1/KanR | 1, 2, S2, S3, S4 |
| MKT179 | J23118 – RBS P22N – T1 | ColE1/KanR | S1 |
| MKT180 | J23118 – RBS BIV-Tat – T1 | ColE1/KanR | S1 |
| MKT182 | J23118 – RBS HIV-REV – T1 | ColE1/KanR | S1 |
| MKT184 | J23118 – RBS λ N – T1 | ColE1/KanR | 1 |
| AB 1.083 | J23118 – RBS λ N-G6 – T1 | ColE1/KanR | 1 |
| JP 2.005 | J23118 – RBS λ N-G8 – T1 | ColE1/KanR | 1, 2 |
| EO 1.036 | J23118 – RBS λ N-G13 – T1 | ColE1/KanR | 1 |
| AB 1.125 | J23118 – RBS λ N-G24 – T1 | ColE1/KanR | 1, S2 |
| AB 1.093 | J23118 – RBS λ N-G28 – T1 | ColE1/KanR | 1 |
| EO 1.037 | J23118 – RBS λ N-G31 – T1 | ColE1/KanR | 1, S3 |
| EO 1.038 | J23118 – RBS λ N-G32 – T1 | ColE1/KanR | 1, S4 |
| EO 1.023 | J23118 – RBS λ N-G24_FH57 – T1 | ColE1/KanR | S2 |
| AB 1.129 | J23118 – RBS λ N-G28_FH5 – T1 | ColE1/KanR | 2 |
| AB 1.130 | J23118 – RBS λ N-G28_FH35 – T1 | ColE1/KanR | 2, 5 |
| EO 1.001 | J23118 – RBS λ N-G28_FH101 – T1 | ColE1/KanR | 2 |
| EO 1.002 | J23118 – RBS λ N-G28_FH108 – T1 | ColE1/KanR | 2 |
| EO 1.005 | J23118 – RBS λ N-G28_FH109 – T1 | ColE1/KanR | 2 |
| EO 1.008 | J23118 – RBS λ N-G28_FH117 – T1 | ColE1/KanR | 2 |
| EO 1.011 | J23118 – RBS λ N-G28_FH133 – T1 | ColE1/KanR | 2 |
| EO 1.018 | J23118 – RBS λ N-G28_FH262 – T1 | ColE1/KanR | 2, 5 |
| EO 1.054 | J23118 – RBS λ N-G31_FH10 – T1 | ColE1/KanR | S3 |
| EO 1.055 | J23118 – RBs λ N-G31_FH20 – T1 | ColE1/KanR | S3 |
| EO 1.056 | J23118 – RBS λ N-G31_FH28 – T1 | ColE1/KanR | S3 |
| EO 1.057 | J23118 – RBS λ N-G31_FH40 – T1 | ColE1/KanR | S3 |
| EO 1.058 | J23118 – RBS λ N-G31_FH47 – T1 | ColE1/KanR | S3 |
| EO 1.059 | J23118 – RBS λ N-G31_FH53 – T1 | ColE1/KanR | S3 |
| EO 1.060 | J23118 – RBS λ N-G31_FH81 – T1 | ColE1/KanR | S3 |
| EO 1.061 | J23118 – RBS λ N-G31_FH174 – T1 | ColE1/KanR | S3 |
| EO 1.062 | J23118 – RBS λ N-G32_FH8 – T1 | ColE1/KanR | S4 |
| EO 1.063 | J23118 – RBS λ N-G32_FH11 – T1 | ColE1/KanR | S4 |
| EO 1.064 | J23118 – RBS λ N-G32_FH112 – T1 | ColE1/KanR | S4 |
| EO 1.065 | J23118 – RBS λ N-G32_FH152 – T1 | ColE1/KanR | S4 |

| | | | |
|---|---|---|---|
| EO 1.066 | J23118 – RBS $\lambda$ N-G32_FH164 – T1 | ColE1/KanR | S4 |
| MKT249 | LuxR – T1 – pLux – RBS $\lambda$ N – T1 | ColE1/KanR | 3 |
| AB 1.135 | LuxR – T1 – pLux – RBS $\lambda$ N G28_FH35 – T1 | ColE1/KanR | 3 |
| EO 1.067 | LuxR – T1 – pLux – RBS $\lambda$ N G28_FH262 – T1 | ColE1/KanR | 3 |
| MKT780 | LuxR – T1 – pLux – RBS $\lambda$ N – T1 (pRL1342) | RSF1010/CmR | 3 |
| MKT781 | LuxR – T1 – pLux – RBS $\lambda$ N G28_FH262 – T1 (prL1342) | RSF1010/CmR | 3 |
| MKT432 | J23118 – MicF(1-13)SS – T1 | p15A/AmpR | 5 |
| MKT602 | J23118 – obgE-sfgfp – T1 | pSC101/CmR | 5 |
| MKT702 | J23118 – lrp-sfgfp – T1 | pSC101/CmR | 5 |
| MKT591 | J23118 – RybB – T1 | p15A/AmpR | 5 |

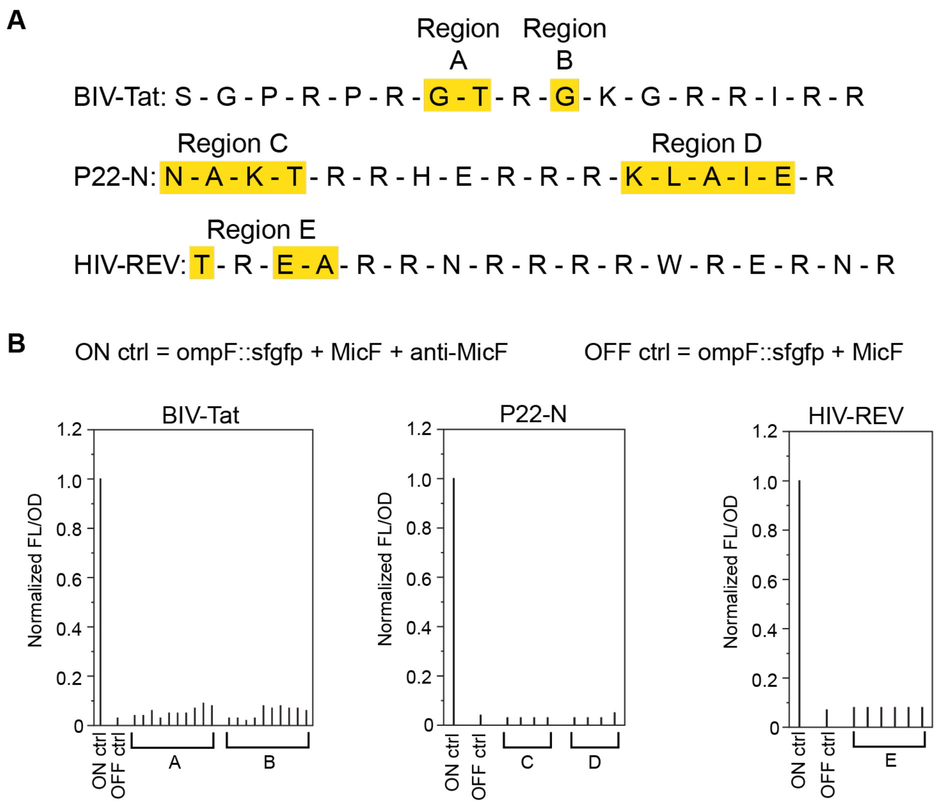


**Figure S1.** Additional randomization results for BIV-Tat, P22-N, and HIV-REV. (A) Amino acid sequences for each ARM with regions randomized highlighted in yellow. (B) Although no colonies appeared to express SFGFP from the visual screen, several individual colonies were selected for growth in liquid culture. FL/OD for each individual culture was normalized to the ON control.

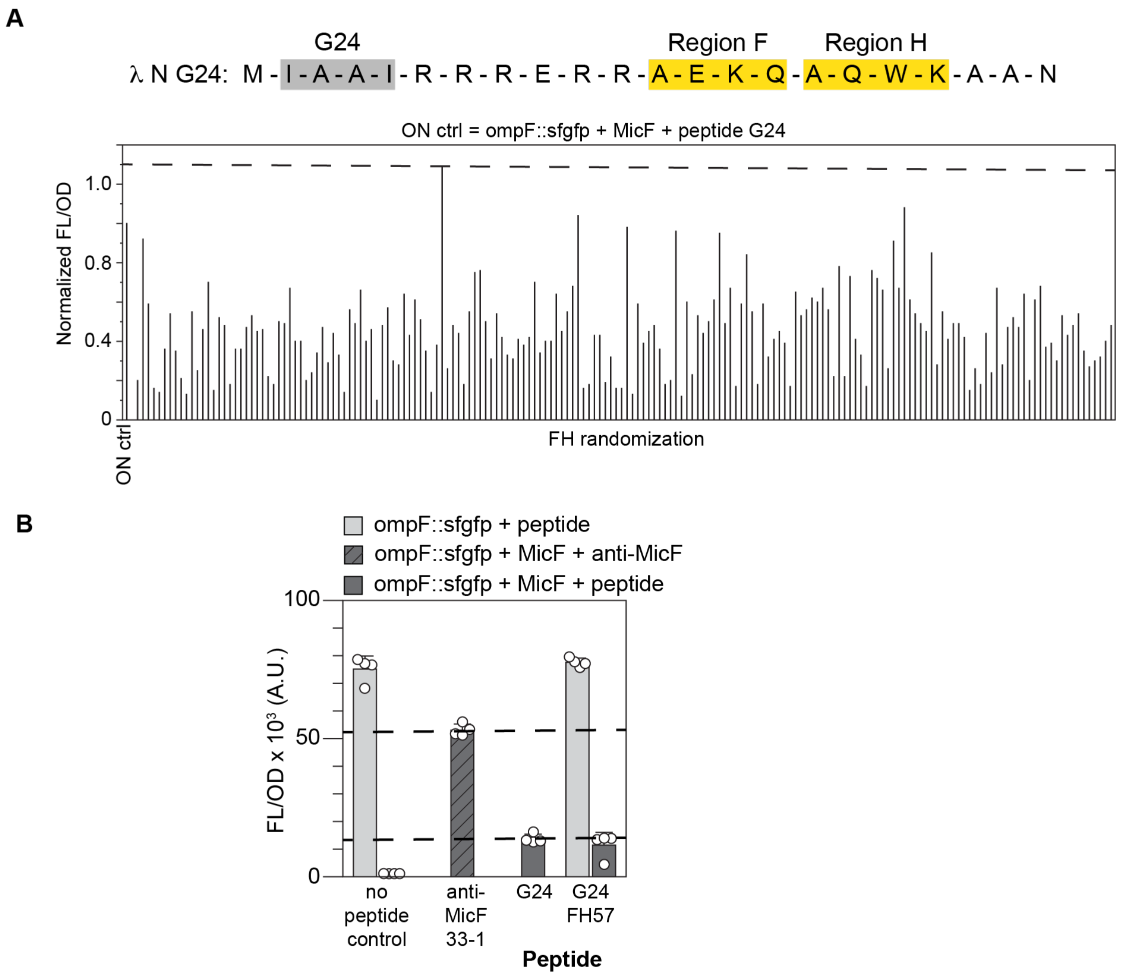


**Figure S2.** Randomization of peptide λ N G24. (A) Randomization screen. The AA sequence of the λ N G24 peptide is shown with the randomized regions F and H highlighted in yellow. Individual colonies from the randomization cloning were grown in liquid culture. FL/OD for each individual culture was normalized to the ON ctrl which was the *ompF::sfgfp* reporter with MicF and the λ N G24 peptide. Peptides above the threshold depicted by the dashed line were sequenced. (B) Peptides above the threshold in (A) with clean, full-length peptide sequences were cloned and tested in *E. coli* along with the *ompF::sfgfp* reporter and MicF. Bars show mean FL/OD. Error bars represent the standard deviations of four biological replicates, shown as open circles. Dashed lines are drawn for comparison at the FL/OD of the anti-MicF control (upper line) and λ N G24 peptide (lower line).

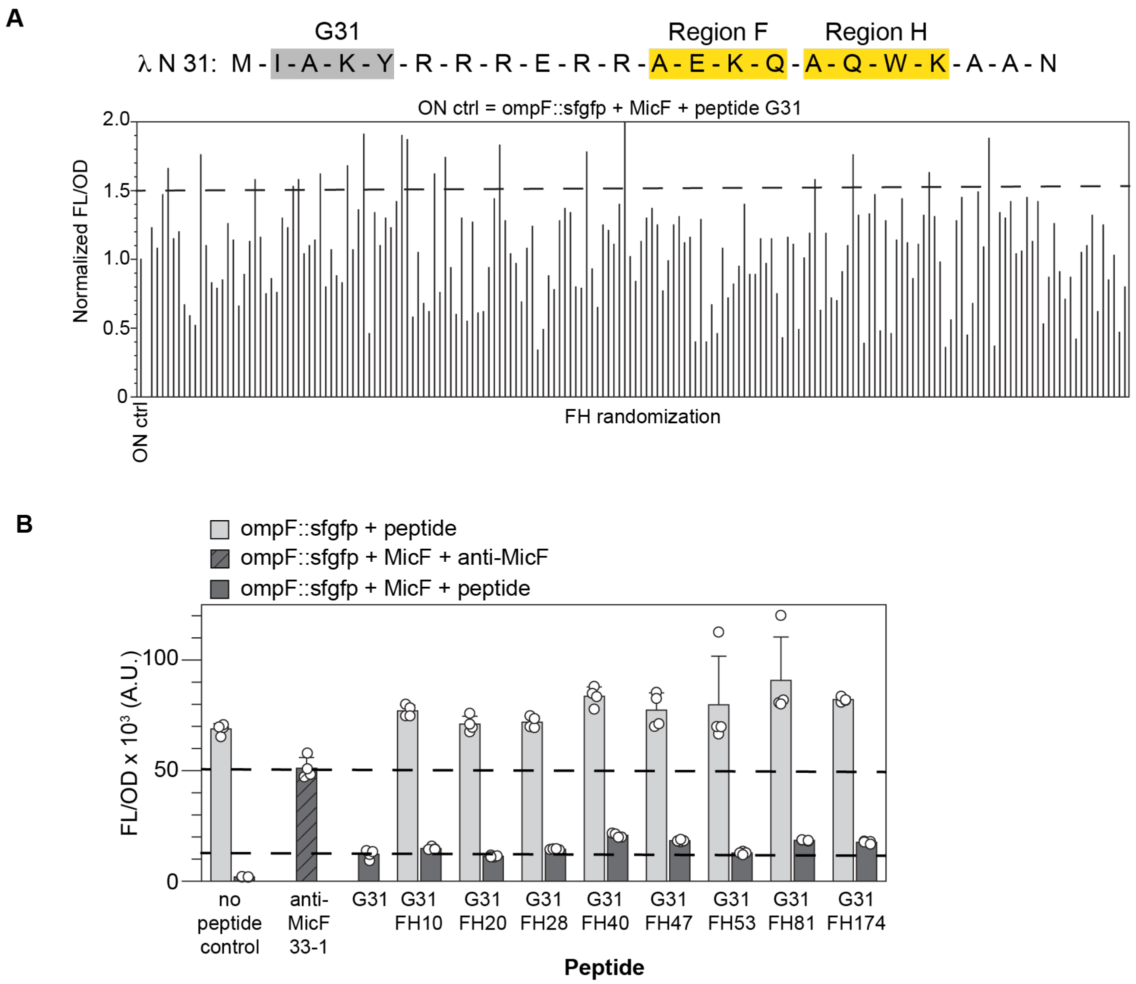


**Figure S3.** Randomization of peptide λ N G31. (A) Randomization screen. The AA sequence of the λ N G31 peptide is shown with the randomized regions F and H highlighted in yellow. Individual colonies from the randomization cloning were grown in liquid culture. FL/OD for each individual culture was normalized to the ON ctrl which was the *ompF::sfgfp* reporter with MicF and the λ N G31 peptide. Peptides above the threshold depicted by the dashed line were sequenced. (B) Peptides above the threshold in (A) with clean, full-length peptide sequences were cloned and tested in *E. coli* along with the *ompF::sfgfp* reporter and MicF. Bars show mean FL/OD. Error bars represent the standard deviations of four biological replicates, shown as open circles. Dashed lines are drawn for comparison at the FL/OD of the anti-MicF control (upper line) and λ N G31 peptide (lower line).

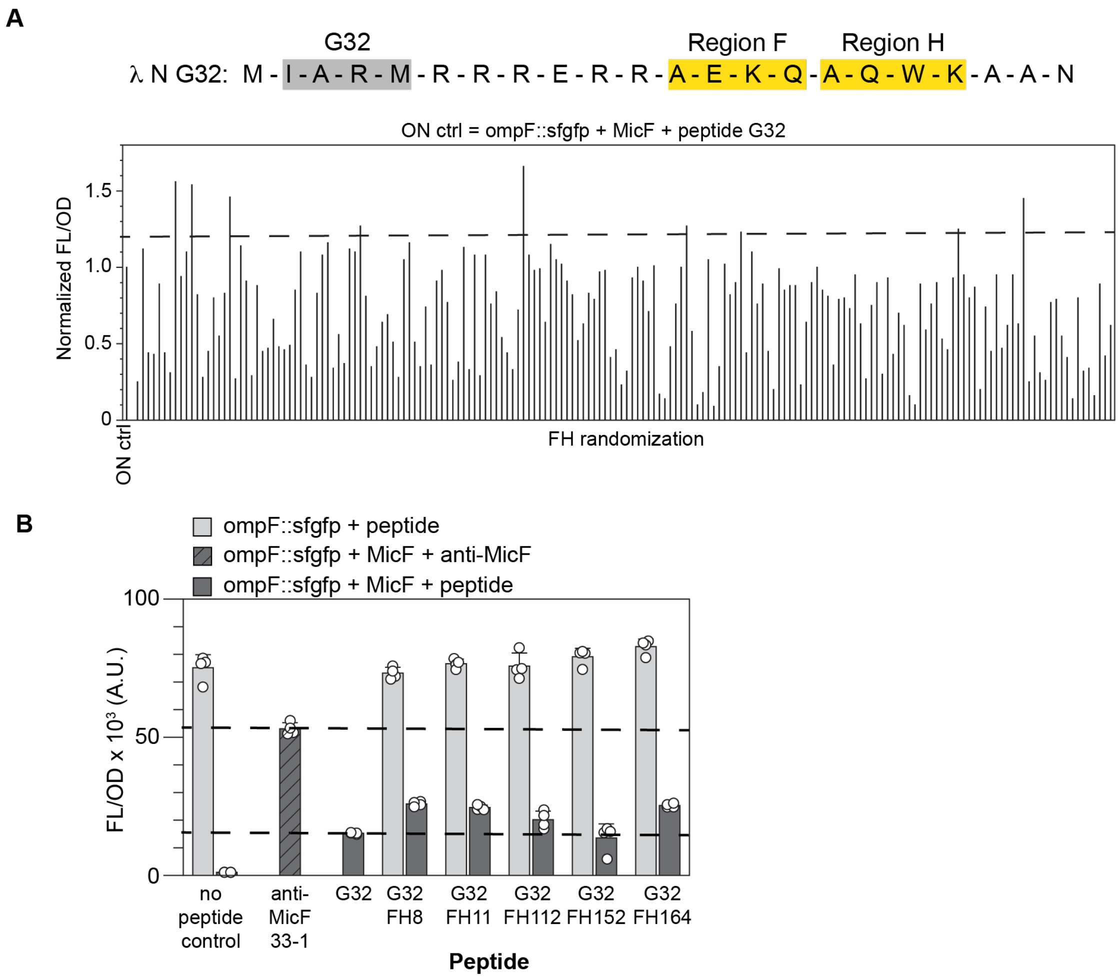


**Figure S4.** Randomization of peptide λ N G32. (A) Randomization screen. The AA sequence of the λ N G32 peptide is shown with the randomized regions F and H highlighted in yellow. Individual colonies from the randomization cloning were grown in liquid culture. FL/OD for each individual culture was normalized to the ON ctrl which was the *ompF::sfgfp* reporter with MicF and the λ N G32 peptide. Peptides above the threshold depicted by the dashed line were sequenced. (B) Peptides above the threshold in (A) with clean, full-length peptide sequences were cloned and tested in *E. coli* along with the *ompF::sfgfp* reporter and MicF. Bars show mean FL/OD. Error bars represent the standard deviations of four biological replicates, shown as open circles. Dashed lines are drawn for comparison at the FL/OD of the anti-MicF control (upper line) and λ N G32 peptide (lower line).

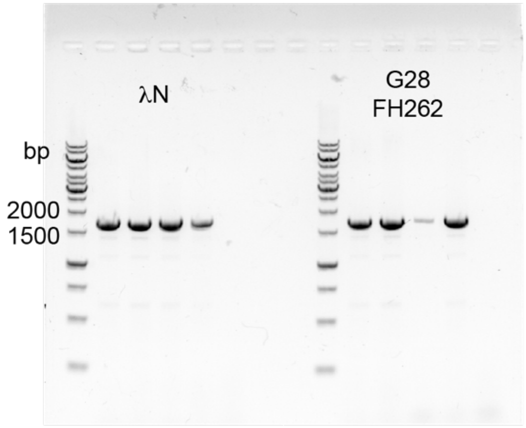


**Figure S5.** PCR to confirm conjugation of peptide carrying plasmid. Colony PCR was performed on four independent colonies from *E. coli* MG1655 that received the λ N and G28_FH262 peptides via conjugation. The GeneRuler 1kb DNA ladder (ThermoFisher Scientific SM0311) was used for size comparison.

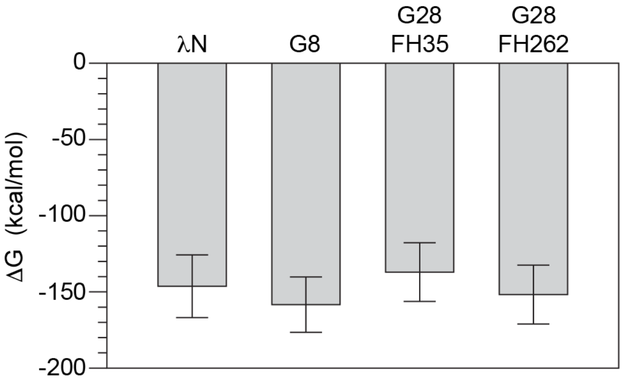


**Figure S6.** Free energy analysis of peptide-MicF complexes at 10,000 frames. Analyses were performed using the Molecular Mechanics/Poisson-Boltzmann Surface Area method. Bars represent averaged energies of 1,000 frames sampled across the 10,000-frame simulation of each complex. Error bars represent the standard deviation of the 1,000 conformations used for the analyses.

**Table S5.** Electrostatic energy contribution per peptide residue in λ N-MicF MD simulation from MMPBSA analysis.

| Peptide Residue | Electrostatic Energy (kcal/mol) |
|---|---|
| MET (M) 1 | -49.0 |
| ASP (D) 2 | 391.7 |
| ALA (A) 3 | -17.2 |
| GLN (Q) 4 | -12.5 |
| THR (T) 5 | -8.9 |
| ARG (R) 6 | -446.5 |
| ARG (R) 7 | -460.3 |
| ARG (R) 8 | -399.6 |
| GLU (E) 9 | 413.1 |
| ARG (R) 10 | -449.4 |
| ARG (R) 11 | -442.5 |
| ALA (A) 12 | -16.9 |
| GLU (E) 13 | 434.9 |
| LYS (K) 14 | -433.3 |
| GLN (Q) 15 | -21.4 |
| ALA (A) 16 | -17.3 |
| GLN (Q) 17 | -15.6 |
| TRP (W) 18 | -7.4 |
| LYS (K) 19 | -442.2 |
| ALA (A) 20 | -17.3 |
| ALA (A) 21 | -17.1 |
| ASN (N) 22 | 100.1 |

**Table S6.** Electrostatic energy contribution per peptide residue in G8-MicF MD simulation from MMPBSA analysis.

| Peptide Residue | Electrostatic Energy (kcal/mol) |
|---|---|
| MET (M) 1 | -54.1 |
| THR (T) 2 | -8.9 |
| ALA (A) 3 | -19.1 |
| ILE (I) 4 | -16.7 |
| ASN (N) 5 | -4.8 |
| ARG (R) 6 | -485.3 |
| ARG (R) 7 | -505.8 |
| ARG (R) 8 | -455.7 |
| GLU (E) 9 | 487.0 |
| ARG (R) 10 | -474.8 |
| ARG (R) 11 | -541.8 |
| ALA (A) 12 | -19.9 |
| GLU (E) 13 | 475.8 |
| LYS (K) 14 | -515.0 |
| GLN (Q) 15 | -30.9 |
| ALA (A) 16 | -18.9 |
| GLN (Q) 17 | -15.3 |
| TRP (W) 18 | -1.0 |
| LYS (K) 19 | -507.7 |
| ALA (A) 20 | -17.8 |
| ALA (A) 21 | -17.4 |
| ASN (N) 22 | 104.2 |

**Table S7.** Electrostatic energy contribution per peptide residue in G28_FH35-MicF MD simulation from MMPBSA analysis.

| Peptide residue | Electrostatic Energy (kcal/mol) |
|---|---|
| MET (M) 1 | -55.4 |
| ILE (I) 2 | -17.6 |
| ALA (A) 3 | -21.3 |
| LYS (K) 4 | -513.6 |
| HIS (H) 5 | -17.1 |
| ARG (R) 6 | -500.1 |
| ARG (R) 7 | -528.5 |
| ARG (R) 8 | -499.9 |
| GLU (E) 9 | 492.6 |
| ARG (R) 10 | -475.9 |
| ARG (R) 11 | -455.6 |
| ILE (I) 12 | -16.2 |
| LEU (L) 13 | -12.7 |
| PHE (F) 14 | -9.1 |
| TYR (Y) 15 | -13.8 |
| HIS (H) 16 | -13.3 |
| ILE (I) 17 | -14.1 |
| ARG (R) 18 | -378.7 |
| ASP (D) 19 | 387.0 |
| ALA (A) 20 | -14.4 |
| ALA (A) 21 | -13.7 |
| ASN (N) 22 | 79.7 |

**Table S8.** Electrostatic energy contribution per peptide residue in G28_FH262-MicF MD simulation from MMPBSA analysis.

| Peptide residue | Electrostatic Energy (kcal/mol) |
|---|---|
| MET (M) 1 | -62.1 |
| ILE (I) 2 | -17.0 |
| ALA (A) 3 | -19.8 |
| LYS (K) 4 | -517.6 |
| HIS (H) 5 | -20.3 |
| ARG (R) 6 | -493.4 |
| ARG (R) 7 | -517.4 |
| ARG (R) 8 | -505.5 |
| GLU (E) 9 | 467.4 |
| ARG (R) 10 | -483.6 |
| ARG (R) 11 | -517.3 |
| CYS (C) 12 | -18.5 |
| LEU (L) 13 | -11.3 |
| LEU (L) 14 | -13.0 |
| ASN (N) 15 | -8.7 |
| SER (S) 16 | -13.9 |
| LEU (L) 17 | -11.0 |
| LEU (L) 18 | -12.7 |
| CYS (C) 19 | -19.9 |
| ALA (A) 20 | -18.0 |
| ALA (A) 21 | -16.8 |
| ASN (N) 22 | 102.4 |

**Table S9.** AA-nucleotide Contact analysis for λ N-MicF MD simulation.

| Peptide AA residue # | MicF nucleotide residue # | # of interacting AA-nucleotide atom pairs | Fraction of simulation time atom pairs interact |
|---|---|---|---|
| MET (M) 1 | U 3 | 224 | 3.0 |
| MET (M) 1 | C 6 | 291 | 1.9 |
| MET (M) 1 | U 5 | 191 | 1.8 |
| MET (M) 1 | A 10 | 277 | 1.2 |
| MET (M) 1 | C 2 | 147 | 1.1 |
| MET (M) 1 | U 93 | 189 | 1.1 |
| ASP (D) 2 | U 88 | 68 | 5.8 |
| ASP (D) 2 | U 89 | 143 | 4.8 |
| ALA (A) 3 | U 89 | 96 | 16.2 |
| ALA (A) 3 | U 88 | 82 | 12.0 |
| GLN (Q) 4 | U 89 | 176 | 20.7 |
| GLN (Q) 4 | U 92 | 52 | 13.3 |
| GLN (Q) 4 | U 93 | 183 | 10.9 |
| GLN (Q) 4 | C 58 | 86 | 9.9 |
| GLN (Q) 4 | U 90 | 39 | 6.6 |
| ARG (R) 6 | A 65 | 263 | 28.9 |
| ARG (R) 6 | U 88 | 159 | 10.3 |
| ARG (R) 6 | C 64 | 186 | 1.3 |
| ARG (R) 7 | U 63 | 198 | 29.2 |
| ARG (R) 7 | C 58 | 142 | 27.7 |
| ARG (R) 7 | U 93 | 221 | 13.7 |
| ARG (R) 7 | U 61 | 105 | 7.6 |
| ARG (R) 7 | C 64 | 163 | 1.4 |
| ARG (R) 8 | U 91 | 81 | 10.9 |
| ARG (R) 8 | C 58 | 183 | 10.4 |
| ARG (R) 8 | U 92 | 36 | 1.8 |
| GLU (E) 9 | U 3 | 122 | 1.3 |
| ARG (R) 10 | C 64 | 352 | 43.8 |
| ARG (R) 10 | U 63 | 264 | 17.8 |
| ARG (R) 10 | A 65 | 122 | 11.6 |
| ARG (R) 11 | C 58 | 264 | 29.1 |
| ARG (R) 11 | U 63 | 157 | 16.0 |
| ARG (R) 11 | C 57 | 82 | 7.2 |
| ARG (R) 11 | U 59 | 124 | 2.9 |

| | | | |
|---|---|---|---|
| LYS (K) 14 | U 63 | 236 | 15.6 |
| LYS (K) 14 | U 62 | 116 | 10.6 |
| LYS (K) 14 | U 59 | 105 | 8.0 |
| GLN (Q) 15 | C 57 | 212 | 21.5 |
| GLN (Q) 15 | C 56 | 154 | 5.9 |
| ALA (A) 16 | U 21 | 142 | 6.1 |
| ALA (A) 16 | U 20 | 137 | 5.6 |
| ALA (A) 16 | A 19 | 126 | 3.7 |
| ALA (A) 16 | U 22 | 122 | 2.7 |
| GLN (Q) 17 | U 21 | 161 | 14.6 |
| GLN (Q) 17 | U 20 | 133 | 10.2 |
| GLN (Q) 17 | U 59 | 112 | 9.7 |
| GLN (Q) 17 | A 60 | 64 | 3.1 |
| TRP (W) 18 | U 59 | 264 | 37.9 |
| TRP (W) 18 | C 58 | 139 | 11.4 |
| TRP (W) 18 | C 57 | 242 | 10.7 |
| TRP (W) 18 | C 56 | 71 | 3.7 |
| LYS (K) 19 | A 23 | 349 | 10.3 |
| LYS (K) 19 | C 55 | 145 | 9.8 |
| LYS (K) 19 | C 56 | 222 | 9.3 |
| LYS (K) 19 | U 22 | 325 | 9.3 |
| LYS (K) 19 | U 21 | 348 | 8.0 |
| LYS (K) 19 | U 20 | 218 | 5.6 |
| LYS (K) 19 | C 57 | 203 | 2.6 |
| ALA (A) 20 | U 22 | 157 | 9.6 |
| ALA (A) 20 | U 21 | 141 | 6.7 |
| ALA (A) 20 | U 20 | 132 | 3.2 |
| ALA (A) 21 | U 59 | 110 | 11.2 |
| ASN (N) 22 | U 59 | 195 | 4.1 |

**Table S10.** AA-nucleotide Contact analysis for G8-MicF MD simulation.

| Peptide AA residue # | MicF nucleotide residue # | # of interacting AA-nucleotide atom pairs | Fraction of simulation time atom pairs interact |
|---|---|---|---|
| MET (M) 1 | A 60 | 110 | 2.8 |
| MET (M) 1 | U 93 | 145 | 2.6 |
| MET (M) 1 | U 59 | 130 | 1.8 |
| THR (T) 2 | C 58 | 82 | 3.6 |
| ILE (I) 4 | U 92 | 27 | 9.9 |
| ASN (N) 5 | U 91 | 68 | 9.8 |
| ASN (N) 5 | U 89 | 10 | 3.7 |
| ARG (R) 6 | C 58 | 64 | 12.8 |
| ARG (R) 6 | C 58 | 146 | 10.5 |
| ARG (R) 6 | U 92 | 90 | 10.4 |
| ARG (R) 6 | A 60 | 44 | 6.2 |
| ARG (R) 6 | C 58 | 60 | 3.4 |
| ARG (R) 7 | C 58 | 54 | 10.0 |
| ARG (R) 7 | U 92 | 71 | 2.3 |
| ARG (R) 8 | U 93 | 133 | 31.5 |
| ARG (R) 8 | U 59 | 43 | 8.3 |
| ARG (R) 8 | U 63 | 21 | 8.3 |
| GLU (E) 9 | U 93 | 76 | 7.2 |
| ARG (R) 10 | C 58 | 59 | 4.9 |
| ARG (R) 10 | C 64 | 46 | 1.6 |
| ARG (R) 11 | U 93 | 88 | 24.6 |
| ARG (R) 11 | U 59 | 47 | 9.1 |
| ARG (R) 11 | C 57 | 28 | 5.4 |
| ALA (A) 12 | U 93 | 31 | 5.5 |
| LYS (K) 14 | U 89 | 47 | 6.5 |
| LYS (K) 14 | A 60 | 26 | 4.3 |
| LYS (K) 14 | U 90 | 25 | 3.5 |
| GLN (Q) 15 | U 63 | 61 | 16.0 |
| GLN (Q) 15 | U 90 | 29 | 9.5 |
| GLN (Q) 15 | U 89 | 36 | 7.1 |
| GLN (Q) 15 | C 57 | 59 | 4.3 |
| TRP (W) 18 | C 64 | 152 | 48.3 |
| TRP (W) 18 | C 64 | 44 | 3.0 |
| LYS (K) 19 | U 91 | 67 | 2.5 |
| ASN (N) 22 | G 1 | 131 | 1.3 |

**Table S11.** AA-nucleotide Contact analysis for G28_FH35-MicF MD simulation.

| Peptide AA residue # | MicF nucleotide residue # | # of interacting AA-nucleotide atom pairs | Fraction of simulation time atom pairs interact |
|---|---|---|---|
| MET (M) 1 | U 90 | 81 | 2.7 |
| MET (M) 1 | U 91 | 51 | 1.3 |
| ILE (I) 2 | U 90 | 63 | 3.6 |
| ILE (I) 2 | U 24 | 98 | 2.5 |
| ILE (I) 2 | U 91 | 65 | 2.5 |
| ALA (A) 3 | U 51 | 18 | 1.1 |
| LYS (K) 4 | C 55 | 29 | 1.2 |
| HIS (H) 5 | U 91 | 64 | 3.5 |
| ARG (R) 6 | U 52 | 97 | 7.8 |
| ARG (R) 6 | U 51 | 31 | 1.9 |
| ARG (R) 6 | U 62 | 120 | 1.1 |
| ARG (R) 7 | C 57 | 137 | 10.4 |
| ARG (R) 7 | C 56 | 131 | 8.4 |
| ARG (R) 7 | A 60 | 60 | 5.7 |
| ARG (R) 7 | U 61 | 37 | 2.3 |
| ARG (R) 8 | A 60 | 65 | 10.2 |
| ARG (R) 8 | U 52 | 76 | 4.0 |
| GLU (E) 9 | A 60 | 8 | 1.0 |
| ARG (R)10 | A 60 | 107 | 16.8 |
| ARG (R)10 | U 61 | 141 | 10.7 |
| ARG (R)10 | U 62 | 80 | 6.5 |
| ARG (R)10 | U 63 | 61 | 2.4 |
| ARG (R)11 | A 60 | 173 | 25.3 |
| ARG (R)11 | U 59 | 80 | 5.4 |
| PHE (F) 14 | A 60 | 57 | 16.4 |

**Table S12.** AA-nucleotide Contact analysis for G28_FH262-MicF MD simulation.

| Peptide AA residue # | MicF nucleotide residue # | # of interacting AA-nucleotide atom pairs | Fraction of simulation time atom pairs interact |
|---|---|---|---|
| MET (M) 1 | U 52 | 302 | 7.5 |
| MET (M) 1 | C 55 | 263 | 6.2 |
| MET (M) 1 | C 58 | 147 | 5.0 |
| MET (M) 1 | U 53 | 264 | 4.2 |
| MET (M) 1 | A 54 | 133 | 1.5 |
| ILE (I) 2 | U 90 | 181 | 9.3 |
| ILE (I) 2 | U 93 | 172 | 6.1 |
| ILE (I) 2 | A 54 | 136 | 1.6 |
| ILE (I) 2 | U 92 | 111 | 1.5 |
| ALA (A) 3 | U 92 | 61 | 11.4 |
| ALA (A) 3 | U 93 | 27 | 2.0 |
| ALA (A) 3 | C 56 | 17 | 1.4 |
| ALA (A) 3 | C 55 | 37 | 1.1 |
| LYS (K) 4 | U 92 | 47 | 7.9 |
| LYS (K) 4 | U 89 | 81 | 6.0 |
| LYS (K) 4 | C 58 | 89 | 4.9 |
| LYS (K) 4 | U 90 | 39 | 1.3 |
| HIS (H) 5 | U 92 | 134 | 22.2 |
| HIS (H) 5 | U 91 | 135 | 16.9 |
| HIS (H) 5 | C 64 | 15 | 4.3 |
| ARG (R) 6 | U 91 | 203 | 32.1 |
| ARG (R) 6 | U 89 | 222 | 14.3 |
| ARG (R) 6 | U 90 | 57 | 8.8 |
| ARG (R) 6 | U 88 | 208 | 8.5 |
| ARG (R) 6 | C 64 | 28 | 4.8 |
| ARG (R) 7 | A 65 | 245 | 26.6 |
| ARG (R) 7 | C 64 | 170 | 25.8 |
| ARG (R) 7 | U 88 | 115 | 5.3 |
| ARG (R) 7 | U 89 | 46 | 4.7 |
| ARG (R) 8 | C 58 | 141 | 13.2 |
| ARG (R) 8 | C 56 | 48 | 7.9 |
| ARG (R) 8 | C 57 | 21 | 6.6 |
| ARG (R) 8 | C 64 | 17 | 5.2 |
| ARG (R) 8 | U 59 | 66 | 1.5 |

| | | | |
|---|---|---|---|
| GLU (E) 9 | C 64 | 67 | 14.8 |
| ARG (R)10 | C 64 | 242 | 28.3 |
| ARG (R)10 | U 63 | 251 | 13.4 |
| ARG (R)10 | A 65 | 43 | 1.0 |
| ARG (R)11 | U 63 | 238 | 31.0 |
| ARG (R)11 | U 59 | 198 | 28.1 |
| ARG (R)11 | C 64 | 54 | 11.4 |
| ARG (R)11 | A 60 | 192 | 8.0 |
| LEU (L)13 | U 63 | 124 | 2.6 |
| LEU (L)13 | U 62 | 122 | 2.4 |
| LEU (L)14 | U 63 | 170 | 17.6 |
| LEU (L)14 | U 62 | 180 | 11.8 |
| LEU (L) 18 | C 57 | 69 | 2.1 |
| LEU (L) 18 | U 59 | 57 | 1.4 |
| CYS (C) 19 | U 92 | 33 | 6.2 |
| ALA (A) 20 | U 92 | 37 | 1.2 |
| ALA (A) 21 | U 92 | 87 | 6.0 |
| ALA (A) 21 | U 93 | 88 | 2.6 |
